# Conserved role of EGFR signaling in apoptotic cell recognition and processing across different phagocytes

**DOI:** 10.64898/2026.06.10.731277

**Authors:** Laura Filomena Comi, Simon Berger, Ambra Villani, Michael Daube, Francesca Peri, Alex Hajnal, Silvan Spiri

**Affiliations:** Department of Molecular Life Science, University of Zürich, Winterthurerstrasse 190, 8057 Zürich; PhD program in Molecular Life Sciences, University of Zürich; present address: Department of Molecular and Cell Biology, University of California, Berkeley, CA, 94720, USA

## Abstract

Efferocytosis is an essential process that clears dying cells from tissues and prevents inflammation. However, the mechanisms by which apoptotic cells recruit and activate phagocytes remain poorly understood. Here, we show that EGFR signaling is both necessary and sufficient for efficient efferocytosis in two diverse phagocytes: the gonadal sheath cells of *C. elegans* and the zebrafish microglia. In *C. elegans*, loss of LET-23 EGFR or its downstream effector MPK-1 ERK in the sheath cells impairs the recognition and degradation of apoptotic germ cells. Germ cells undergoing apoptosis secrete the EGF ligand LIN-3, which promotes their recognition and engulfment. Overexpression of EGF in non-apoptotic germ cells is sufficient to trigger their engulfment, indicating that EGF can function as an engulfment signal independently of apoptotic cell death. In zebrafish larvae, pharmacological inhibition of EGFR signaling reduces microglial motility, recognition of apoptotic neurons, and corpse clearance in the optic tectum. Similar to *C. elegans,* an ectopic source of EGF serves as an attractive cue that promotes microglial recruitment. Our findings suggest that EGFR signaling controls an evolutionarily conserved efferocytosis module that coordinates the recognition and processing of apoptotic corpses in different types of phagocytes.

## Introduction

Programmed cell death (apoptosis) is an essential process in multicellular organisms, removing unnecessary or damaged cells to support healthy development, maintain tissue homeostasis, and prevent disease^1–3^. However, the elimination of dying cells starts, rather than ends, with apoptosis. After cells initiate apoptosis, phagocytes must efficiently recognize and degrade them to ensure their elimination from the tissue via a process termed efferocytosis. In vertebrates, this task is performed largely by tissue-resident macrophages, including microglia in the brain. However, both invertebrates and vertebrates also use non-professional phagocytes, such as epithelial cells and fibroblasts, for local clearance of apoptotic cells^1,4,5^. Defective efferocytosis, such as failures in apoptotic-cell recognition or degradation by phagocytes, contributes to numerous chronic inflammatory conditions, autoimmune diseases, and neurodegenerative disorders^6,7^. Consequently, efferocytosis occurs across biological scales and throughout the animal kingdom. Many advances in understanding this process have been made using model organisms^1,8,9^.

Apoptotic cell clearance is a multi-step process that requires phagocytes to coordinate several distinct functions. First, phagocytes must detect apoptotic cells from a distance and migrate towards them in a chemotactic response to "find-me" signals^10^. Second, phagocytes must establish contact with the apoptotic cell via surface receptors that recognize "eat-me" signals, such as phosphatidylserine (PS) exposed on the apoptotic cell membrane^11^. Third, activation of phagocytic receptors triggers signaling cascades that promote cytoskeletal remodeling required for the formation of a phagosome, a process termed engulfment^12^. Finally, the apoptotic cell corpse undergoes digestion through sequential fusion with endosomal and lysosomal vesicles, a process termed phagosome maturation^13,14^. While many genes controlling these individual steps have been identified, how cells coordinate them to achieve efficient corpse clearance remains poorly understood. Moreover, different phagocyte populations rely on distinct, context-dependent receptor repertoires^15,16^. In particular, it is unclear whether the mechanisms are conserved, as efferocytosis occurs in evolutionarily distant organisms and involves both professional and non-professional phagocytes. Moreover, phagocytes can increase their phagocytic capacity to adapt to elevated levels of cell death^17^. Yet it is unclear how a high phagocytic capacity is sustained over long periods of elevated apoptosis, for example, during multiple waves of neuronal cell death in the maturing brain.

Another underexplored aspect of efferocytosis is how intercellular signaling mediated by classical growth factors and their receptors contributes to the clearance of apoptotic cells. Receptor tyrosine kinases such as the epidermal growth factor receptor (EGFR) are best known for their roles in cell proliferation, survival, and differentiation^18^. EGFR dysregulation has been extensively studied in cancer and developmental disorders^19,20^. However, potential functions of EGFR signaling in phagocytosis have received little attention. Yet, growth factor pathways are well-positioned to integrate information about tissue state: EGF family ligands can be produced by stressed or dying cells^21^, and EGFR signaling can modulate cell motility, cytoskeletal dynamics, and vesicular trafficking^22^, all of which are essential processes during efferocytosis. Therefore, a key question in the field is whether apoptotic cells use growth factor signaling, particularly EGF ligands, to recruit neighboring phagocytes to clear apoptotic corpses.

To address these questions, we investigated two in vivo models, in which a small number of phagocytes clear many apoptotic cells during a phase of sustained physiological cell death: The *C. elegans* gonadal sheath cells (**Figure 1A**), which are non-professional phagocytes that remove apoptotic germ cells, and the zebrafish larval microglia, which are professional phagocytes that clear apoptotic neurons in the developing optic tectum (**Figure 4A**)^23^. Unlike developmental cell death in *C. elegans*, which follows an invariant cell lineage where the same 131 somatic cells undergo apoptosis^24^, more than half of the developing oocytes in the *C. elegans* hermaphrodite germline undergo stochastic apoptosis, a process termed physiological germ cell death^25,26^. These apoptotic cells are cleared exclusively by the germline-encasing, epithelial-like sheath cells from the somatic gonad. This comparative approach allowed us to ask whether a conserved signaling mechanism coordinates efferocytosis across evolutionarily distant phagocytes.

**Figure 1.**
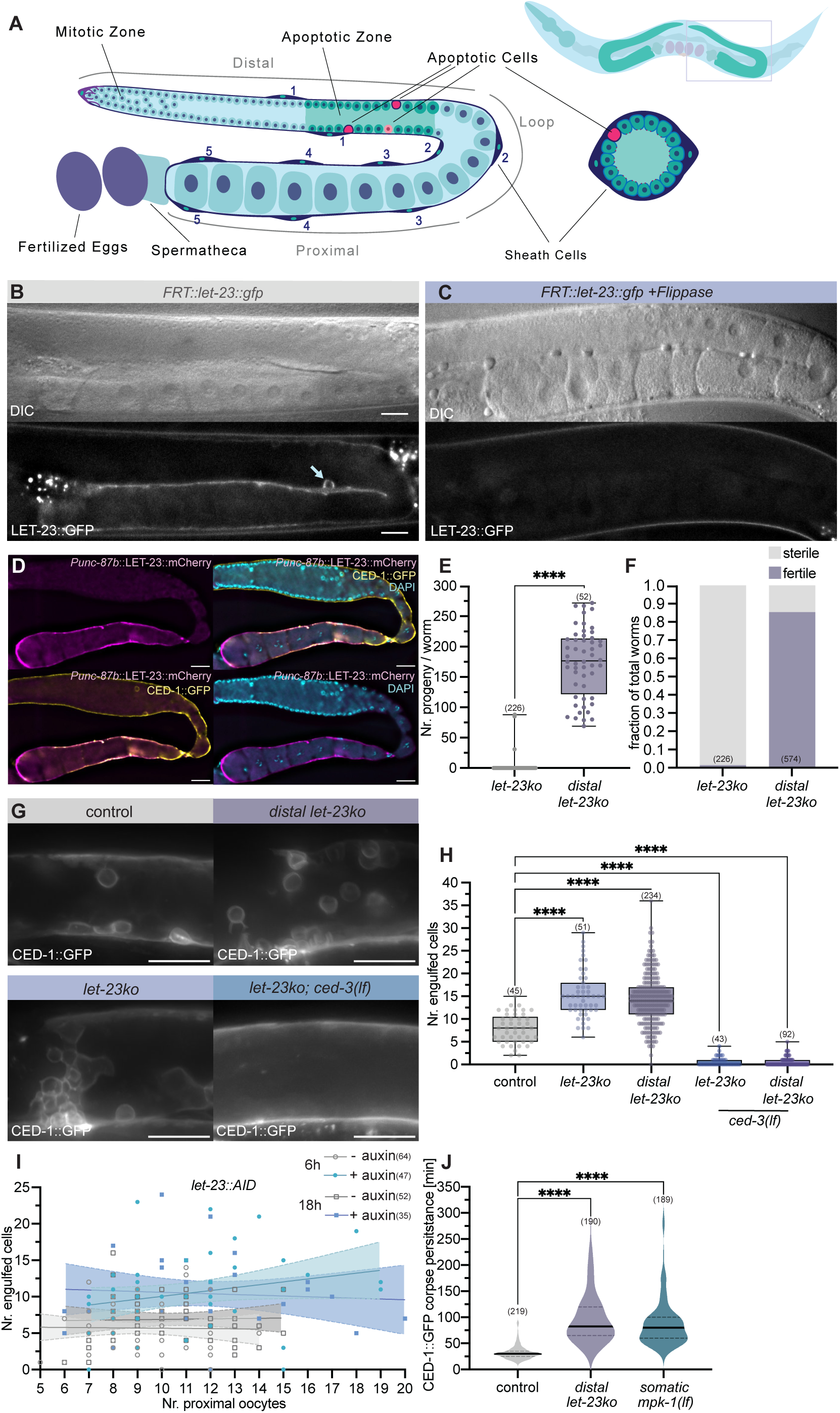
*let-23 egfr* in the *C. elegans* sheath cells positively regulates apoptotic germ cell engulfment. **(A)** Schematic of one gonad arm in an adult *C. elegans* hermaphrodite. Left: longitudinal view, right: cross-section in the apoptotic zone. Apoptotic cells are colored in red, and the phagocytic sheath cells encasing the germline are in dark blue. Distal (1 & 2) and proximal (3-5) sheath cell pairs are indicated. **(B)** DIC (top) and GFP channel (bottom) of an animal carrying the endogenously *gfp*-and *FRT*-tagged conditional *let-23* allele (*FRT::let-23::gfp*). A LET-23::GFP-positive encircled germ cell corpse is indicated by the arrow. **(C)** DIC (top) and *let-23::gfp* expression (bottom) in an animal carrying the endogenously *gfp*-and *FRT*-tagged *let-23* allele crossed with the somatic gonad-specific Flippase *Pckb-3::FLP* (*let-23ko*). The LET-23::GFP signal in the sheath cells and surrounding the apoptotic germ cells is lost. Note the ovulation defect visible as broken-down proximal oocytes in the DIC image. **(D)** Dissected and DAPI-stained gonad of a *distal let-23ko* animal carrying the proximal sheath cell-specific *Punc-87B::let-23::mCherry* rescue transgene and the *ced-1::gfp* reporter. LET-23::mCherry (magenta) is expressed in the proximal sheath cells, and CED-1::GFP (yellow) is expressed in both proximal and distal sheath cells. A single row of proximal oocytes is visible in the DAPI channel (cyan), indicating restored ovulation. **(E)** Brood size and **(F)** fertility analysis of the *let-23ko* and *distal let-23ko* strains. **(G)** Maximum intensity projections of CED-1::GFP in the apoptotic zone of indicated strains. Whole-gonad CED-1::GFP expression with corresponding DIC images is shown in **Figure S1C**. **(H)** Numbers of CED-1::GFP-positive germ cell corpses in the indicated strains. For the *distal let-23ko* and *distal let-23ko; ced-3(lf)* strains, three independent transgenic lines were analyzed to control for variability in extrachromosomal arrays. The complete data are shown in **Figure S1D**. **(I)** Quantification of *let-23::AID* animals treated with auxin for 6h and 18h and untreated controls. Quantification of engulfed cells for the 6h and 18h auxin-treatment timepoints are shown in **Figure S1E**. The number of proximal oocytes was used as a proxy for the severity of the ovulation defect and plotted against the number of CED-1::GFP-positive germ cell corpses. Severe ovulation defects (number of proximal oocytes too large to count) were excluded. Solid lines indicate the results of a linear regression with 95% confidence intervals (shaded area). Data points assigned an oocyte number of 20 were not included in the linear regression analysis. **(J)** Persistence of CED-1::GFP-positive germ cell corpses in the indicated strains. Young adult worms were imaged in 5 min intervals for 8 h. Only CED-1::GFP-positive cells, which were degraded during the recording, are included. Six animals were recorded for each strain. All worms analyzed for this figure were staged at 24 hpL4, except for **(E)**, where offspring were counted during the first 4 days of adulthood, and **(J)**, where young adults were imaged for 8h. The numbers in brackets in each graph refer to the number of animals scored, except for **(J)**, where the numbers of tracked cells are indicated. Boxplots show the median and interquartile range, and the whiskers show the minimum and maximum values. Violin plots with smoothing show the median (black line) and quartiles (dotted lines). Statistical tests used: **(E)** unpaired, two-tailed Mann-Whitney test, **(H)** Kruskal-Wallis test with Dunn’s multiple comparisons correction, **(J)** Brown-Forsythe and Welch test with Dunnett’s T3 multiple comparisons correction. Statistical significance is indicated as ns (p ≥ 0.05), * (p < 0.05), ** (p < 0.01), *** (p < 0.001), **** (p < 0.0001). Scale bars: 20 µm.

Here, we show that EGFR signaling coordinates multiple steps of apoptotic cell clearance in both models. In *C. elegans*, apoptotic germ cells express the EGF ligand LIN-3, which activates LET-23 EGFR/RAS/MAPK signaling in the sheath cells to promote corpse recognition, engulfment, and degradation. Ectopic LIN-3 expression in the germline is sufficient to induce engulfment of non-apoptotic germ cells. In zebrafish larvae, EGFR inhibition reduces microglial motility, recognition of apoptotic neurons and degradation of engulfed material, whereas an ectopic source of EGF attracts microglia. Together, these findings identify EGFR signaling as a conserved mechanism that coordinates phagocyte recruitment with the uptake and processing of apoptotic cells across diverse phagocyte types.

## Results

### Loss of LET-23 EGFR in the gonadal sheath cells results in the accumulation of apoptotic germ cells

While analyzing the expression pattern of an endogenously *gfp*- and *FRT*- tagged *let-23 egfr* allele ^27^ in *C. elegans* hermaphrodites, we observed LET-23::GFP protein expression in the gonadal sheath cells, specifically enriched around apoptotic germ cells in the distal gonads (**Figure 1B**). To investigate a potential role of *let-23* in germ cell efferocytosis, we combined the *FRT::let-23::gfp* allele with a somatic gonad-specific flippase transgene, driven by the *ckb-3* promoter (termed *let-23ko* hereafter)^28^. *let-23ko* resulted in loss of the LET-23::GFP signal in the sheath cells, a penetrant ovulation defect, and complete sterility (**Figure 1C,E,F**). Both phenotypes have previously been reported for *let-23(lf)* alleles^29^. To investigate the role of *let-23* in efferocytosis independently from its function during ovulation, we took advantage of the spatial segregation of the two processes. Germ cell efferocytosis is mostly carried out by the sheath cell pairs 1 and 2, which encase the apoptotic zone in the distal germline. On the other hand, ovulation requires myoepithelial contraction of the proximal sheath cell pairs 3, 4 and 5. By expressing a *let-23::mCherry* rescue construct under the proximal sheath cell-specific *unc-87b* promoter (**Figure S1A,B**)^30^, we were able to rescue the ovulation defect and restore fertility in most *let-23ko* animals (**Figure 1D-F**).

To test whether loss of *let-23* in distal sheath cells affects efferocytosis, we used a *ced-1::gfp* reporter^31,32^, which is enriched in sheath cells surrounding engulfed apoptotic corpses. One-day-old adult *let-23ko* hermaphrodites (24 hours post L4 stage; 24hpL4) contained elevated numbers of apoptotic corpses in the late pachytene zone compared to wild-type controls, as did *let-23ko* animals expressing *Punc-87b::let-23::mCherry* in the proximal gonads (termed *distal let-23ko*, **Figure 1** **G,H**). Furthermore, *let-23ko* animals carrying the *ced-3(n717) caspase lf* mutation, which blocks apoptosis, lacked engulfed germ cells, indicating that the accumulation of engulfed cells results from physiological germ cell death rather than necrotic death or other causes^25,33^ (**Figure 1** **G,H**).

To rule out that defects in somatic gonad development indirectly cause elevated numbers of engulfed cells in the *let-23ko* and *distal let-23ko* mutants, we created an endogenously *degron*-tagged *let-23* allele (*let-23::AID*), allowing us to downregulate LET-23 protein in adults using the auxin-inducible degradation system^34,35^. *let-23::AID* animals carrying a pan-somatic *tir-1* driver and the *ced-1::gfp* reporter were transferred to auxin-containing plates at 6 or 18 hours post-L4, when somatic gonad development was complete. Auxin-induced LET-23 degradation (*let-23kd*) at 6h or 18h post L4 resulted in a significant increase in the numbers of engulfed germ cells at 24hpL4 compared to *let-23::AID* control animals grown without auxin (**Figure S1E**). Thus, the accumulation of apoptotic corpses does not originate from a loss of *let-23* function during larval development. Moreover, a linear regression of the number of engulfed cells against the number of proximal oocytes indicated that the ovulation and engulfment defects in *let-23kd* animals are not co-dependent. *let-23kd* animals (+ auxin) had a higher number of engulfed cells than control animals (- auxin) containing the same number of oocytes at both time points (**Figure 1I**; i.e., the intercepts of control and *let-23kd* are significantly different (p<0.0001), but the slopes are not (p=0.1866 for 6h, p=0.6689 for 18h)).

Next, we aimed to determine whether the engulfment phenotype was due to an elevated rate of germ cell death or delayed corpse degradation, as both defects would increase the number of engulfed germ cells. Engulfed germ cells remain mostly stationary and are surrounded by CED-1::GFP for an extended time before decreasing in size and GFP intensity until they are no longer detectable. This allows corpse persistence to be quantified by measuring the duration of CED-1::GFP labeling. Using microfluidic long-term imaging devices^36^, we measured CED-1::GFP persistence via time-lapse imaging. Engulfed corpses persisted, on average, more than three times longer in *distal let-23ko* animals than in wild-type controls, suggesting that the accumulation of engulfed germ cells is caused by delayed corpse degradation rather than enhanced germ cell death (**Figure 1J**).

Taken together, these results show that loss of LET-23 EGFR function in distal sheath cells leads to delayed degradation of apoptotic germ cells. This phenotype is not due to defects in gonad development and uncovers a role of LET-23 EGFR in the distal somatic gonad that is independent of the ovulation defects caused by loss of *let-23* in the proximal somatic gonad.

### EGFR promotes corpse clearing by activating MPK-1 ERK signaling in the sheath cells

Next, we tested whether LET-23 EGFR activates the ERK signaling cascade in sheath cells to promote the degradation of apoptotic corpses. The *C. elegans* ERK ortholog MPK-1 is required for various aspects of germline apoptosis^25^. In particular, loss of the germline-specific MPK-1B isoform (**Figure 2A**) almost completely blocks germ cell apoptosis^26^. Interestingly, we previously reported that a short (4-hour) incubation at the restrictive temperature of 25 °C in a temperature-sensitive *mpk-1(ga111ts)* allele^37^ results in a slightly increased, rather than a decreased, number of engulfed germ cells^26^. However, combining the *mpk-1(ts)* allele with a somatic- or sheath cell-specific *mpk-1* rescue construct resulted in significantly reduced numbers of engulfed cells even below wild-type levels^38^. This suggests that *mpk-1* functions in the somatic sheath cells during the clearance of apoptotic germ cells^26^.

**Figure 2.**
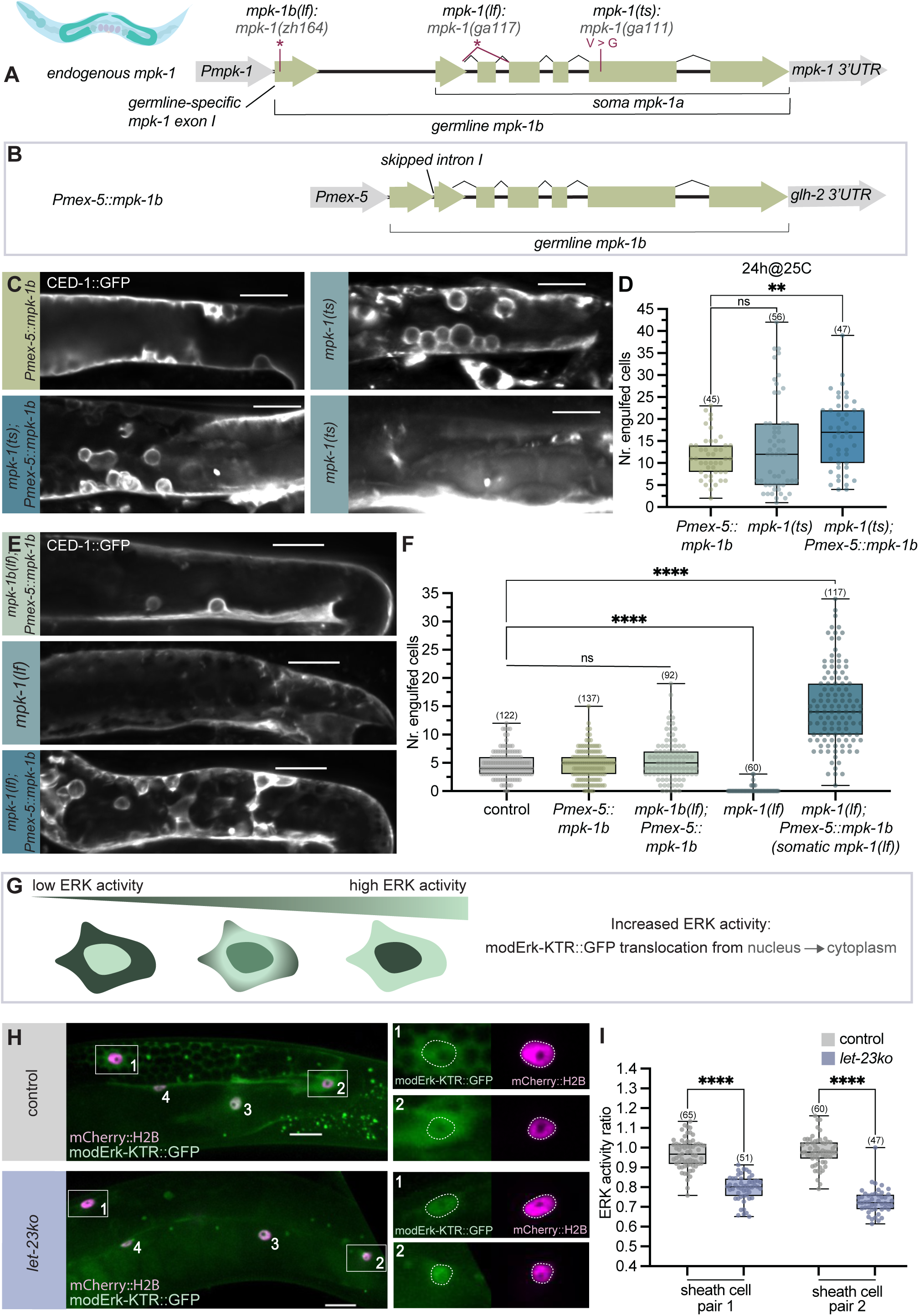
*mpk-1 erk* signaling in the *C. elegans* sheath cells is necessary for efficient corpse degradation. **(A)** Structure of the *mpk-1* locus encoding germline-specific *mpk-1b* and somatic *mpk-1a* isoforms. The three *mpk-1* alleles used are shown. Red asterisks indicate premature stop codons. *mpk-1(ga117)* causes skipping of the 3^rd^ exon, leading to a premature stop codon. **(B)** Structure of germline-specific *mpk-1b* transgene (*Pmex-5::mpk-1b*). **(C)** Maximum intensity projections of CED-1::GFP expression in the apoptotic zone of indicated *mpk-1(ts)* strains. Two examples of *mpk-1(ts)* are shown to illustrate the phenotypic variability. **(D)** Quantification of CED-1::GFP-positive engulfed germ cell corpses in indicated *mpk-1(ts)* strains. **(E)** Maximum intensity projections of CED-1::GFP expression in the apoptotic zone of indicated *mpk-1(lf)* strains. **(F)** Quantification of CED-1::GFP-positive engulfed germ cell corpses in the indicated strains. CED-1::GFP expression in the additional strains analyzed is shown in **Figure S2C**. **(G)** Function of the modERK-KTR biosensor. Increased ERK activity leads to nuclear export of modERK-KTR::GFP. **(H)** modERK-KTR::GFP expression (green) in the sheath cells of wild-type control and *let-23ko* animals. The nuclear mCherry::H2B marker is shown in magenta. Left panels show merged gonad images, highlighting one sheath cell per pair (numbered 1-4). Right panels show magnified views of individual channels in sheath cells 1 and 2. Nuclei are outlined with a dashed circle. **(I)** Quantification of modERK-KTR biosensor activity in sheath cell pairs 1 and 2 of wild-type control and *let-23ko* animals. (See methods for details on the quantification.) All worms analyzed for this figure were staged at 24hpL4. The numbers in brackets in each graph refer to the number of animals scored. Boxplots show the median and interquartile range; whiskers show minimum and maximum values. Statistical tests used: **(D, I)** Brown-Forsythe and Welch test with Dunnett’s T3 multiple comparisons correction, **(F)** Kruskal-Wallis test with Dunn’s multiple comparisons correction. Statistical significance is indicated as ns (p ≥ 0.05), * (p < 0.05), ** (p < 0.01), *** (p < 0.001), **** (p < 0.0001). Scale bars: 20 µm.

Here, we again observed a slight but statistically insignificant increase in corpse numbers in *mpk-1(ts)* single mutants compared to the control, with large animal-to-animal variability (**Figure 2C,D**; variances of the control and *mpk-1(ts)* are significantly different, p<0.0001). To decipher the role of somatic *mpk-1* signaling in germ cell efferocytosis, we introduced a germline-specific *mpk-1* rescue construct by expressing the *mpk-1b* isoform from a germline-specific *mex-5* promoter in the *mpk-1(ts)* background (*Pmex-5::mpk-1b*, **Figure 2B**). The *Pmex-5::mpk-1b* transgene partially rescued the sterility of *mpk-1(ts)* mutants grown at the restrictive temperature (**Figure S2A**). The number of engulfed germ cells in *mpk-1(ts)* mutants with the *Pmex-5::mpk-1b* transgene was significantly increased compared to animals carrying the *Pmex-5::mpk-1b* transgene in a wild-type background (**Figure 2C,D**; see **Figure S2B** for animals grown at 20°C and different durations at 25°C). Thus, restoring *mpk-1* activity in the germ cells of *mpk-1(ts)* mutants results in an accumulation of engulfed germ cells.

Next, we introduced the *Pmex-5::mpk-1b* transgene into *mpk-1* null (*ga117*) mutants^37^ and, as a control, the germline-specific isoform *mpk-1b(zh164)* mutants, both of which contain almost no germ cell corpses and are completely sterile^17^. Germline-specific *mpk-1b* mutants carrying the *Pmex-5::mpk-1b* transgene did not contain more corpses, demonstrating that the *Pmex-5::mpk-1b* transgene is not responsible for the increased number of apoptotic corpses (**Figure 2E,F**, **Figure S2C**). In contrast, *mpk-1* null mutants with the *Pmex-5::mpk-1b* transgene (*somatic mpk-1(lf)*) showed a significant increase in the number of engulfed corpses compared to wild-type controls (**Figure 2E,F, Figure S2C**). Thus, restoring *mpk-1b* activity in the germ cells of *mpk-1* null mutants unmasks the role of *mpk-1* in apoptotic corpse clearance. Moreover, time-lapse imaging of *somatic mpk-1(lf)* mutants revealed an increased lifetime of CED-1::GFP-positive corpses, indicating that the elevated number of engulfed germ cells in *somatic mpk-1(lf)* animals results from a delay in corpse degradation (**Figure 1J**). Together with the *mpk-1(ts)* mutant analysis described above, quantification of *somatic mpk-1(lf)* mutants supports the conclusion that MPK-1 signaling in somatic cells promotes apoptotic corpse clearance.

To test if *let-23 egfr* directly activates *mpk-1* signaling in the sheath cells, we created a sheath cell-specific ERK kinase translocation (ERK-KTR) biosensor to quantify MPK-1 activity (**Figure 2G**). For this purpose, we used a modified version of the original ERK-KTR sensor^39^ (modERK-KTR), which minimizes phosphorylation of the biosensor by Cdk-dependent kinases^40^. We first expressed the modERK-KTR biosensor in the vulval precursor cells (VPCs) and compared its activity with that of the original ERK-KTR biosensor^39^. The modERK-KTR biosensor showed a similar activity pattern as the original ERK-KTR biosensor, with the strongest activity in the primary VPC P6.p and weaker activity in the adjacent VPCs (**Figure S3**). We then created a sheath cell-specific version by expressing modERK-KTR under the control of the *lim-7* enhancer/promoter, which is active in the sheath cell pairs 1 to 4^41^. Highest modERK-KTR activity was detected in sheath cell pairs 1 and 2 (**Figure 2H,I, Figure S4A,B**). To test whether the modERK-KTR biosensor is sensitive to RAS/MAPK signaling in sheath cells, we treated adult worms for 4 hours with the MEK inhibitor U0126. In MEKi-treated animals, modERK-KTR biosensor activity was significantly reduced in all sheath cells (**Figure S4C**). Finally, we introduced the modERK-KTR biosensor into the *let-23ko* strain and found a strong reduction in modERK-KTR biosensor activity in all sheath cells (**Figure 2H,I, Figure S4A,B**) Taken together, these results show that activation of the MPK-1 ERK pathway by LET-23 EGFR in the somatic sheath cells promotes the clearance of apoptotic germ cell corpses.

### LIN-3 EGF expression in germ cells promotes apoptotic corpse recognition and degradation

Next, we aimed to find the source of the EGF ligand that activates LET-23 EGFR signaling in the sheath cells. Three EGF ligands have so far been described in *C. elegans*^42–44^. The canonical EGF ligand LIN-3 controls many cell fate decisions during larval development, while SISS-1 and IGEG-2 regulate predominantly neuronal functions during sleep. Since *lin-3* was previously shown to be expressed in the gonads^45^, we examined endogenous *lin-3* expression using a translational *mNeongreen::lin-3*^46^ reporter and a bicistronic transcriptional reporter consisting of an *SL2::mCherry::H2B* cassette inserted into the *lin-3* locus after the last exon, labeling the nuclei of cells expressing *lin-3* mRNA (**Figure S5A,B**). The transcriptional LIN-3::SL2::mCherry::H2B reporter was expressed in all germ cells (**Figure 3A**). Apoptotic germ cells showed a slight increase in *lin-3* transcription. However, the increased signal intensity could also be due to nuclear compaction or pH changes in the apoptotic corpses. Moreover, we observed faint expression of the translational mNeongreen::LIN-3 reporter around apoptotic cells in 19 of 50 animals (**Figure 3B,C**).

**Figure 3.**
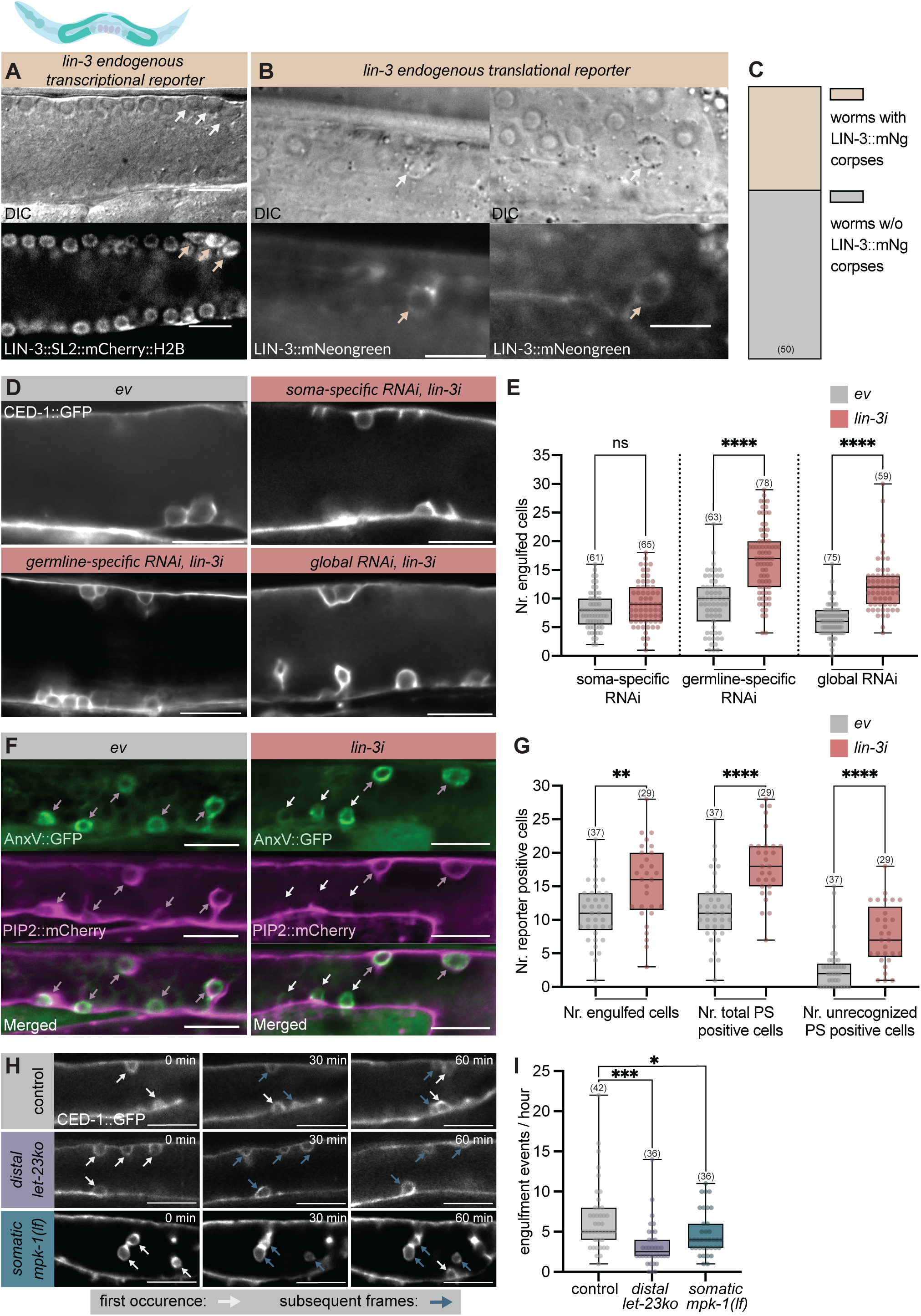
*lin-3 egf* expression in *C. elegans* germ cells promotes apoptotic corpse recognition. **(A)** Endogenous transcriptional LIN-3::SL2::HIS-58::mCherry reporter expression (bottom) in late pachytene-stage germ cells with the corresponding DIC image (top). Arrows point to apoptotic germ cells. **(B)** Endogenous translational mNeongreen::LIN-3 reporter expression. Two examples of LIN-3::mNeongreen-positive apoptotic germ cell corpses are shown (arrows). **(C)** Frequency of LIN-3::mNeongreen-positive corpses. **(D)** CED-1::GFP expression in single z-slices of the apoptotic zone in *lin-3* RNAi-treated (*lin-3i*) worms in global, soma- and germline-specific RNAi strains. (See methods for the tissue-specific RNAi strains used). The empty vector RNAi control (*ev*) is shown for global RNAi. Worms were fed for 24h with *ev* or *lin-3i* bacteria. **(E)** Quantification of CED-1::GFP-positive engulfed germ cell corpses in indicated RNAi strains. **(F)** Expression of AnxV::GFP (green) and PIP2::mCherry (magenta) in *ev* global *lin-3i* animals. Purple arrows indicate AnxV::GFP- and PIP2::mCherry-double positive cells, white arrows indicate AnxV::GFP-positive, PIP2::mCherry-negative cells (i.e., unrecognized PS-positive cells). Worms were treated for 24 h with *ev* or *lin-3i* bacteria. For heat shock-induced AnxV::GFP expression, worms were heat-shocked for 45 min at 33°C and allowed to recover at 20°C for 1.5 h before imaging. **(G)** Quantification of PIP2::mCherry-positive engulfed germ cell corpses (Nr. engulfed cells), total AnxV::GFP positive cells (Nr. PS-positive cells) and AnxV::GFP-positive, PIP2::mCherry-negative cells (Nr. of unrecognized PS-positive cells) in *ev* and global *lin-3i* treated animals. **(H)** CED-1::GFP expression in single z-slices of the apoptotic zone of long-term imaging of indicated strains at three timepoints with 30-minute intervals. CED-1::GFP-positive corpses are indicated with a white arrow at their first appearance and blue arrows in subsequent timepoints. **(I)** Number of successful engulfment events in indicated strains. The first 6 hours after the first corpse occurrence were analyzed as described in methods. Number of animals analyzed: control N = 7, *distal let-23ko* N = 6, *somatic mpk-1(lf)* N= 7. All worms shown in this figure were staged at 24hpL4, except for **(H,I)**, where young adults were imaged for 12 h. The numbers in brackets in the graphs in **(C,E,G)** refer to the number of animals scored and in **(I)** to number of hours where engulfment events were observed. Boxplots show median and interquartile range, whiskers indicate minimum and maximum values. Statistical tests used: **(E,G,I)** Brown-Forsythe and Welch test with Dunnett’s T3 multiple comparisons correction. Statistical significance is indicated as ns (p ≥ 0.05), * (p < 0.05), ** (p < 0.01), *** (p < 0.001), **** (p < 0.0001). Scale bars: 20 µm.

Furthermore, we used tissue-specific *lin-3* RNAi (*lin-3i*) to compare the effects of *lin-3* knockdown in the soma versus the germline. Global *lin-3i* and germline-specific *lin-3i* treatment beginning in the L4 stage until the first day of adulthood (24hpL4) increased the number of engulfed cells, comparable to *let-23ko* and *somatic mpk-1(lf)* mutants. In contrast, soma-specific *lin-3i* resulted only in a slight, non-significant increase in the number of engulfed cells(**Figure 3D,E**). These results suggest that *lin-3* expressed in germ cells positively regulates efferocytosis.

The LIN-3 expression pattern and germline-specific *lin-3i* experiment raised the question of whether LIN-3 may also play a role in corpse recognition before engulfment. The CED-1::GFP reporter we used marks only apoptotic cells that have already been recognized by the sheath cells. To detect unrecognized apoptotic germ cells, we combined a PIP2::mCherry engulfment reporter^26^ with a heat shock-inducible AnnexinV::GFP reporter (AnxV::GFP)^47^. AnnexinV binds to the “eat-me” signal phosphatidyl serine (PS), which is exposed on the outer plasma membrane of cells that initiate apoptosis, already before their engulfment. Combining AnxV::GFP with the PIP2::mCherry engulfment reporter enabled us to distinguish between unrecognized and recognized apoptotic germ cells. Since AnnexinV binding could mask exposed PS and thus interfere with corpse engulfment, we first performed a control experiment comparing engulfment levels in the presence and absence of the AnxV::GFP reporter under the heat-shock and recovery conditions used. We did not observe a significant change in the total number of engulfed (i.e., PIP2::mCherry-positive) germ cells in the presence of the AnxV::GFP reporter under experimental conditions (**Figure S5C**). By contrast, global *lin-3i* caused a significant increase in the number of unrecognized (i.e., AnxV::GFP-positive, PIP2::mCherry-negative) apoptotic germ cells (**Figure 3F,G**). Moreover, quantification of the phagosome formation rates by live imaging revealed a decreased frequency of successful engulfment events in *distal let-23ko* and *somatic mpk-1(lf)* mutants (**Figure 3H,I**). In summary, these data suggest that apoptotic germ cells secrete LIN-3 to promote their engulfment by sheath cells. Thus, LIN-3 is required not only for efficient degradation but also for the recognition of apoptotic germ cells.

### EGFR signaling in zebrafish promotes apoptotic cell recognition and degradation by the microglia

After discovering that the *C. elegans* sheath cells, which are non-professional phagocytes, rely on EGF/EGFR signaling for efficient efferocytosis, we investigated whether the EGFR signaling pathway is a conserved regulator of efferocytosis used by professional phagocytes as well. To address this question, we investigated efferocytosis in the developing zebrafish optic tectum (OT), a well-established model in which a small population of approximately 25-30 microglia clears a large number of apoptotic neurons during normal development (**Figure 4A**)^48^.

We performed pharmacological inhibition of the EGFR signaling in 3 dpf zebrafish larvae. The specific EGFR inhibitor Erlotinib has previously been shown to block EGFR signaling in the developing zebrafish brain at 5 dpf^49^ without causing cardiotoxicity^50^, suggesting that this drug could also be suitable for inhibiting the EGFR in the microglia. We treated 3 dpf larvae with 20 µM Erlotinib for 24 hours and visualized microglia using the *fms::mCherry* reporter. To detect apoptotic neurons, Acridine Orange (AO) staining was performed, which marks apoptotic corpses that microglia have phagocytosed, as well as uncollected apoptotic neurons.

First, we verified that the Erlotinib treatment does not perturb development or neuronal apoptosis. Microglial number in the OT was not changed by Erlotinib treatment (**Figure 4B**). To test whether Erlotinib alters the rate of neuronal apoptosis in the OT independent of microglial clearance, we quantified the number of AO-positive cells in *irf8^st^*^95^ (*irf8-/-*) mutants, which lack microglia at this stage^51^. Erlotinib-treated *irf8-/-* mutants did not show increased numbers of AO-positive cells in the OT or the hindbrain (HB), a brain region with less apoptosis (**Figure 4C,D)**. Altering spot-detection parameters or using higher Erlotinib concentrations did not affect this finding (**Figure S6C-E**). Moreover, OT and HB size as well as relative apoptotic cell density in the two brain regions were not altered by Erlotinib treatment (**Figure S6F-I**).

By contrast, the number of uncollected AO-positive apoptotic neurons (i.e., AO spots outside microglia masks) was increased in Erlotinib-treated larvae, pointing to a defect in the recognition or uptake of apoptotic neurons (**Figure 4E,F**). To further dissect this defect, we performed long-term imaging of the microglia in the OT. Time-lapse analysis revealed that Erlotinib treatment reduced the frequency of successful engulfment events, suggesting that EGFR inhibition compromises the uptake of apoptotic neurons (**Figure 4G**). In addition to the reduced engulfment rate, Erlotinib treatment caused an accumulation of apoptotic corpse material inside microglial cells and an increase in microglial volume (**Figure 4E,H, Figure S6A,B**). Since EGFR inhibition reduced the engulfment rate, the accumulation of apoptotic corpse material is unlikely to result from excessive corpse uptake and instead suggests impaired degradation of internalized apoptotic corpses.

To further investigate the mechanism underlying the accumulation of apoptotic material, we used the *Lamp1-mgfp* reporter, which marks phagolysosomes, i.e., phagosomes at a late stage of maturation after fusion with lysosomes^52^. The number and size of Lamp1-positive vesicles, and the total phagolysosomal area per microglia were all increased in the microglia of Erlotinib-treated larvae (**Figure 5A-C, Figure S7**). By tracking phagolysosome size over time through time-lapse imaging, we found that phagolysosomes in Erlotinib-treated zebrafish shrank at a slower rate compared to controls, confirming that the increased number and size of phagolysosomes in microglia are due to delayed corpse degradation rather than increased efferocytosis (**Figure 5D**, **Video S1**).

**Figure 4.**
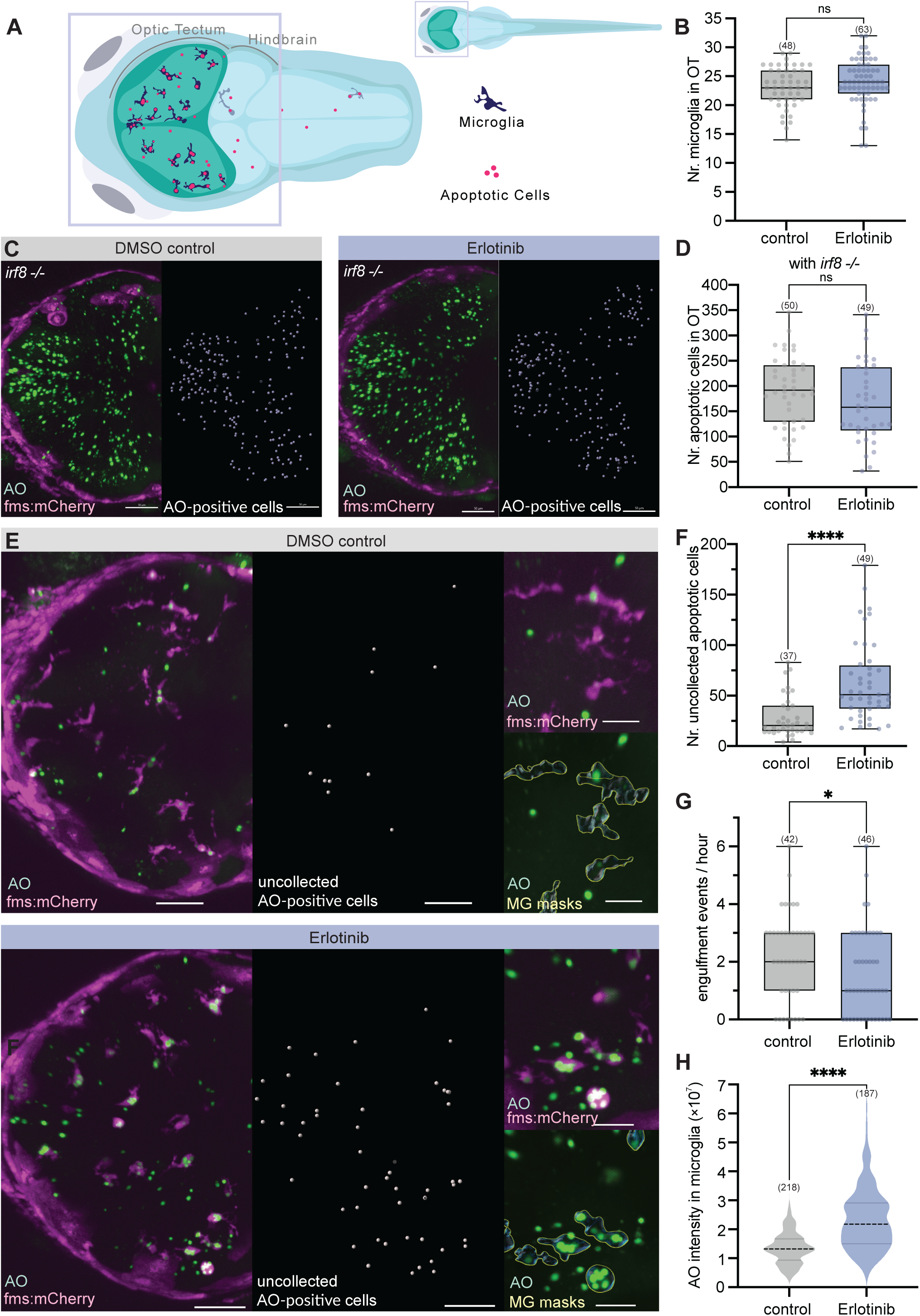
EGFR inhibition in zebrafish OT impairs apoptotic corpse recognition and degradation. **(A)** Schematic overview of the optic tectum (OT) in 3 dpf zebrafish larvae. Microglia are depicted in dark blue, and apoptotic neurons are in pink. **(B)** Total number of microglia in the OT in control and Erlotinib-treated larvae. **(C)** *fms::mCherry* expression (magenta) and AO staining of apoptotic neurons (green) in control and Erlotinib-treated *irf8-/-* mutants (*irf8st95*) that lack microglia in the OT at this stage^51^. Left: Merged maximum intensity projections. Right: AO spot detection as described in methods used to count apoptotic neurons in **(D)**. No *fms::mCherry*-positive microglia were present in the OT of *irf8-/-* mutants, but xanthophores in the skin were labeled with *fms::mCherry*. **(D)** Quantification of the number of apoptotic neurons in control and Erlotinib-treated *irf8-/-* mutants. **(E)** Microglia labeled with *fms::mCherry* (magenta) and apoptotic neurons stained with AO (green) in the OT of control and Erlotinib-treated zebrafish. Left: merged maximum intensity projection. Middle: AO spot detection to count apoptotic neurons outside the microglia (i.e., AO-stained cells outside the microglia masks). See methods for details on the AO spot-detection parameters and microglia masking. Top right: Merged close-up images of *fms::mCherry* expression and AO staining. Bottom right: AO staining merged with the outline of the microglia masks (white lines), used to measure AO intensity within microglia. **(F)** Quantification of uncollected apoptotic neurons (i.e., AO spots outside the microglia masks) in control and Erlotinib-treated larvae. **(G)** Number of successful engulfment events per microglia and hour in three control and Erlotinib-treated larvae each (see methods). **(H)** Quantification of summed AO intensity within microglia in 25 control and 26 Erlotinib-treated larvae (See methods for details on the measurement workflow). The experiments shown in **(B,E-F,H)** and **(C, D)** were conducted simultaneously using the same drug-containing medium to ensure comparability. All zebrafish were staged at 3 dpf. All images shown are maximum-intensity projections, with the top z-slices containing skin xanthophores removed for improved visualization. The numbers in brackets in **(G,H)** refer to the number of microglia analyzed. The numbers in brackets in the graphs in **(B,D,F)** refer to the number of animals analyzed. Boxplots show median and interquartile range, whiskers indicate minimum and maximum values. Violin plots with smoothing in **(H)** show the median (black dotted line) and quartiles (gray lines). Statistical tests used: Unpaired, two-tailed Welch’s tests. Statistical significance is indicated as ns (p ≥ 0.05), * (p < 0.05), ** (p < 0.01), *** (p < 0.001), **** (p < 0.0001). Scale bars: 50 µm, except in the close-ups in **(B):** 20 µm.

**Figure 5.**
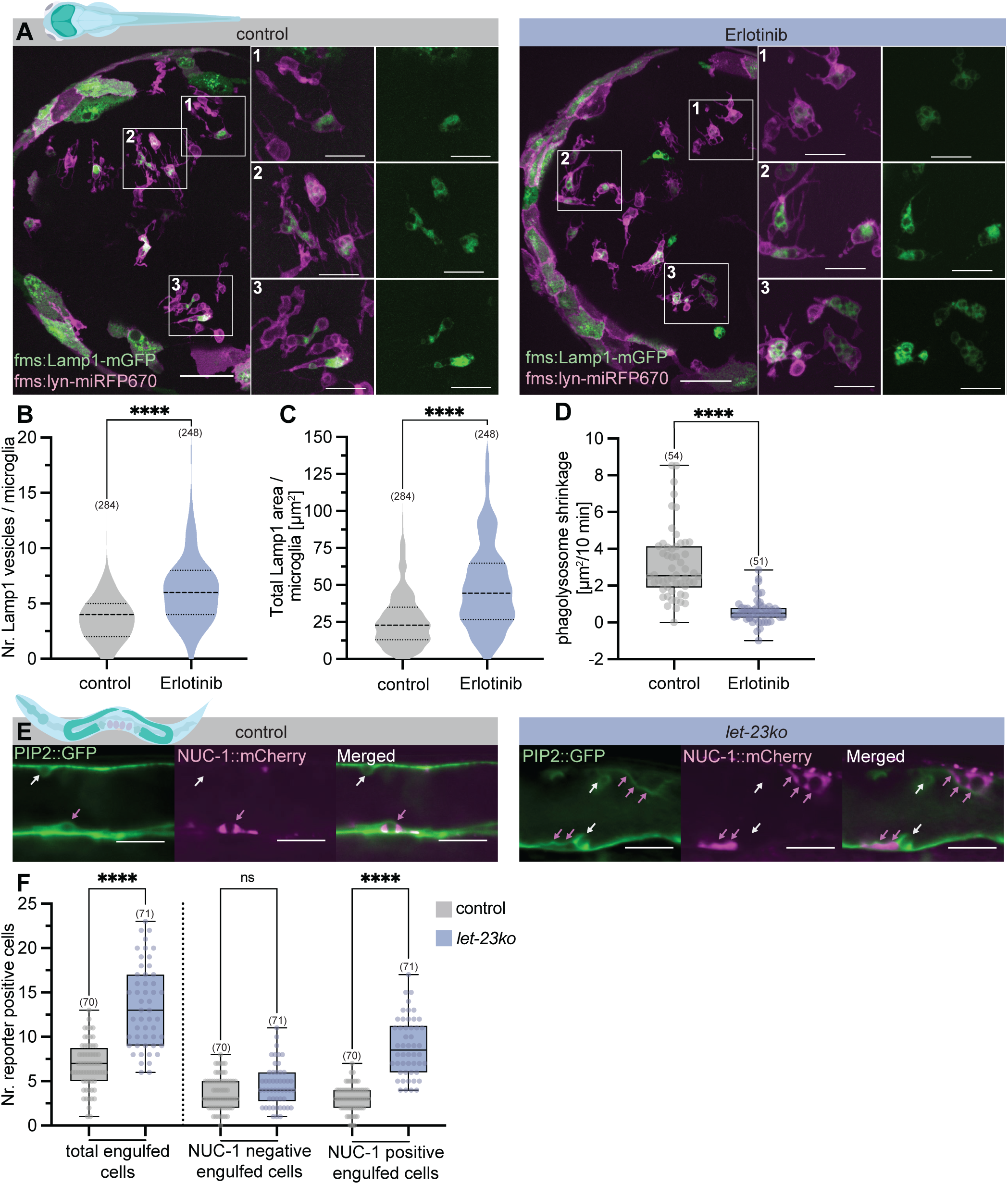
Phagolysosome accumulation and reduced engulfment rate after EGFR inhibition in the zebrafish OT and *C. elegans* sheath cells. **Zebrafish: (A)** Left: Merged maximum intensity projections of *fms::Lamp1-mGFP* (green) expression marking lysosomes and *fms::lyn-miRFP670* expression (magenta) marking microglia membranes in the OT. Right: Magnified view of the three regions outlined with numbered white boxes in the left panels. **(B)** Number of Lamp1-mGFP-positive vesicles per microglia. See methods for measurement parameters. **(C)** Area of Lamp1-mGFP-positive vesicles per microglia. Thirty control and Erlotinib-treated larvae each were analyzed for **(B,C)**. **(D)** Phagolysosomal area shrinkage per 10 minutes measured in three control and Erlotinib-treated larvae each (see methods). All zebrafish larvae were staged at 3 dpf. ***C. elegans*: (E**) Single z-slices showing expression of PIP2::GFP engulfment reporter (green) and NUC-1::mCherry lysosomal marker (magenta) in the germline of control and *let-23ko* animals. White arrows indicate engulfed, NUC-1::mCherry-negative germ cells. Pink arrows indicate engulfed, NUC-1::mCherry-positive cells. **(F)** Number of PIP2::GFP and NUC-1::mCherry reporter-positive cells in control and *let-23ko* animals. Total engulfed cells correspond to all PIP2::GFP-positive cell corpses. All *C. elegans* hermaphrodites were staged at 24hpL4. The numbers in brackets in the graphs in **(B-D)** refer to the number of microglia analyzed, in **(F)** to the number of animals analyzed. Boxplots indicate median and interquartile range, whiskers show minimum and maximum values. Violin plots with smoothing show the median (dashed line) and quartiles (dotted lines). Statistical tests used: **(B-D)** unpaired, two-tailed Welch’s tests; **(F)** Brown-Forsythe and Welch tests with Dunnett’s T3 multiple comparisons correction. Statistical significance is indicated as ns (p ≥ 0.05), * (p < 0.05), ** (p < 0.01), *** (p < 0.001), **** (p < 0.0001). Scale bars: 50 µm in **(A)** left, 20 µm in close-ups in **(A)** and in **(E)**.

These findings in zebrafish larvae prompted us to ask whether the loss of LET-23 EGFR in *C. elegans* sheath cells similarly affects phagolysosomal function. To test this, we combined the PIP2::GFP engulfment reporter with the phagolysosomal marker NUC-1::mCherry^53^. The number of NUC-1::mCherry-positive engulfed (i.e., NUC-1::mCherry and PIP2::GFP double-positive) cells was increased in *let-23ko* animals, but the number of NUC-1::mCherry-negative, engulfed corpses was not significantly changed (**Figure 5E,F)**. Thus, the elevated number of engulfed apoptotic cells in the *let-23ko* strain likely results from the accumulation of germ cell corpses within phagolysosomes. This is consistent with the observation that engulfed corpses persisted more than three times longer in both *distal let-23ko* and *somatic mpk-1(lf)* mutants (**Figure 1J**).

In summary, EGFR inhibition in zebrafish larvae perturbs multiple steps of microglial efferocytosis, from apoptotic-cell uptake to corpse processing in phagolysosomes, without increasing neuronal death or causing obvious developmental defects, consistent with the phagolysosomal degradation defects seen upon loss of *let-23 egfr* in the *C. elegans* sheath cells.

### EGF stimulates microglial mobility in zebrafish and LIN-3 EGF overexpression in C. elegans germ cells causes engulfment

The reduced engulfment rate upon EGFR inhibition in zebrafish suggested a defect in apoptotic cell recognition, prompting us to examine whether EGFR signaling regulates microglial motility. For this purpose, we developed an image analysis pipeline that generates binary masks of individual microglia from time-lapse image stacks and extracts microglial features over time (**Figure 6A**, **Video S2**; additional examples are shown in **Figure S8**; extracted features are illustrated in **Figure S9**. See **Figure S10, Video S3** and **Data S4** for image analysis pipeline details). Using the curated 3D masks, we quantified microglial mobility by calculating frame-to-frame Pearson correlation coefficients. Microglia in Erlotinib-treated larvae exhibited reduced mobility and dynamics (i.e., increased correlation coefficients, **Figure 6C**, **Figure S9I-L**). Tracking individual branch tips over time revealed that both elongation and retraction speeds were decreased by Erlotinib treatment (**Figure 6D**). On the other hand, total branch length and the number of branch tips per microglia cell were unchanged in Erlotinib-treated larvae, suggesting that the reduced mobility was not due to morphological changes (**Figure 6B**, **Figure S9B,D,H**). Only a slight increase in the number of branch trees (i.e., branches with a single root connected to the soma) was observed, indicating a modest reduction in ramification and a shift toward individually rooted branches (**Figure S9C,G**). Thus, when EGFR is inhibited, microglia retain their overall branch morphology but become markedly less motile, which likely underlies the reduced efficiency of engulfment of apoptotic corpses.

**Figure 6.**
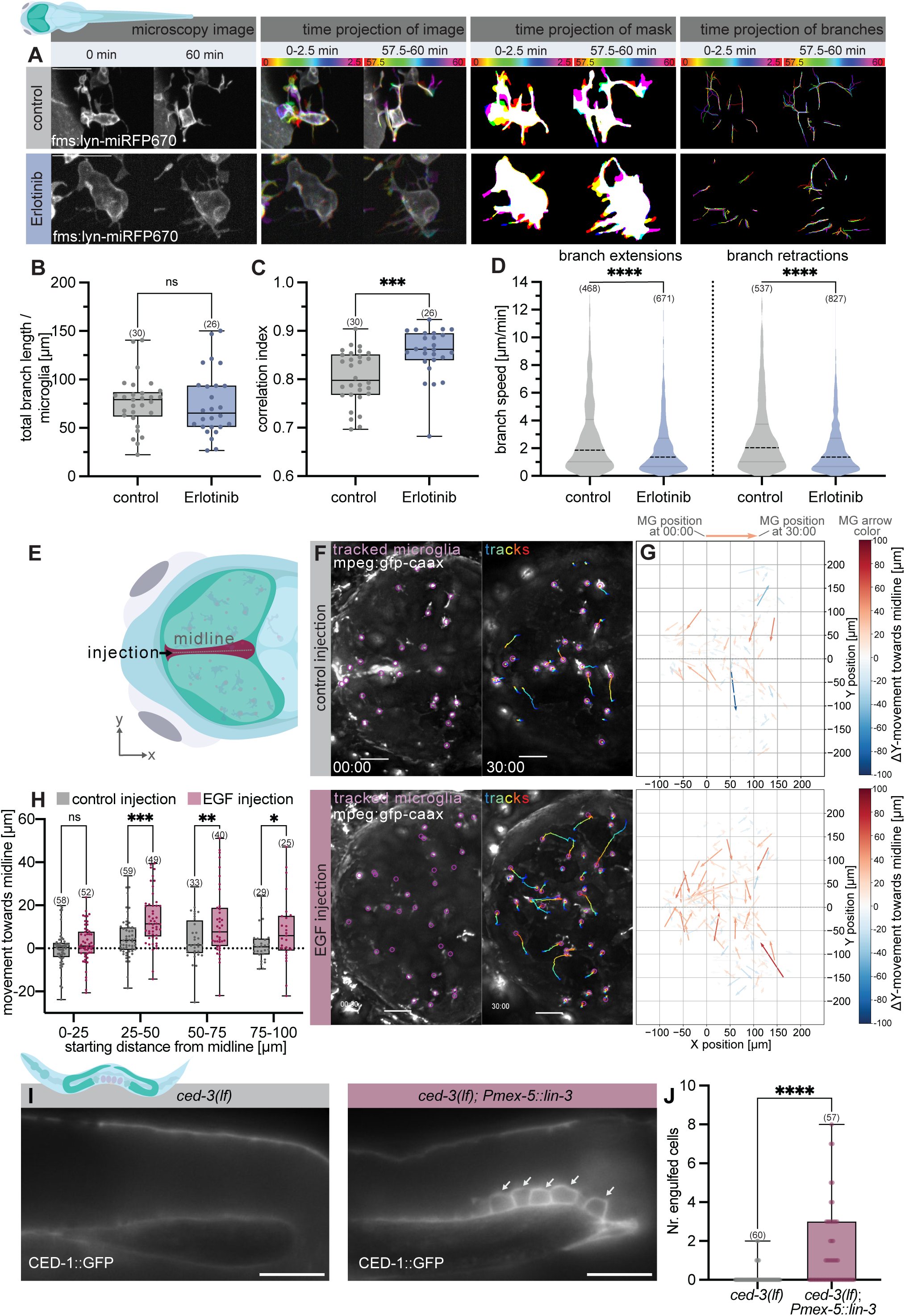
Ectopic EGF is an attractive cue for microglia in the zebrafish OT and an engulfment signal in the *C. elegans* germline. **Zebrafish: (A)** Time-lapse imaging of microglia labeled with *fms::lyn-miRFP670* reporter in 3 dpf larvae. For each condition, the first panel (from left) shows maximum intensity projections of *fms:lyn-miRFP670* signal at the beginning and end of the 60 min recording, the second panel shows color-coded time projections of the raw image, the third panel shows color-coded time projections of the microglia masks, and the fourth panel shows color-coded time projection of microglia branches, each showing projections for five consecutive frames at beginning and end of the recording.. See **Figure S8** for additional examples and **Figure S10** for the image processing workflow. **(B)** Total branch length per microglia in control and Erlotinib-treated larvae. **(C)** Pearson correlation coefficients between consecutive time frames calculated from the microglia masks, as illustrated in **Figure S9I.** Mean correlation coefficients per microglia over 60 min are shown. Additional measurements are shown in **Figure S9J-L**. **(D)** Speed of microglia branch extensions and retractions. Single-tip branches were tracked over time using the root position and angle (see methods and **Figure S10**). **(E)** EGF injections into the midline of the zebrafish OT in 4 dpf larvae. See methods for details. **(F)** Maximum intensity projections of the OT in control and EGF-injected larvae immediately after injection (00:00, left) and at the end of the tracking period (30:00, right). Microglia were labeled with the *mpeg::gfp-caax* membrane marker. The purple dots indicate tracked microglial cell bodies, and the tracks are shown with time-coded color lines. **(G)** Vector plots showing the displacement of microglia over the 30 min tracking period after injections. The combined data of the 8 tracked animals are shown. The arrows are color-coded according to their length and direction (red: movement towards the midline, blue: movement away from midline). In total, 257 microglia were tracked in 8 control and 261 microglia in 8 EGF-injected larvae. **(H)** Distance traveled by microglia towards the midline relative to their starting distance from the midline, grouped in 25 µm intervals. ***C. elegans***: **(I)** Expression of the CED-1::GFP engulfment reporter in the late pachytene zone of (left panel) *ced-3 caspase(lf)* single mutants and (right panel) after germ cell-specific overexpression of *lin-3 egf* in *ced-3 caspase(lf)* background. White arrows point to non-apoptotic, CED-1::GFP-engulfed germ cells. **(J)** Quantification of CED-1::GFP-engulfed germ cells in indicated strains. All *C. elegans* hermaphrodites were staged at 24hpL4. The numbers in brackets in the graphs in **(B,C,H)** refer to the number of microglia analyzed, in **(D)** to the number of tracked microglia branches, and in **(J)** to the number of animals scored. Boxplots show median and interquartile range. Whiskers indicate the minimum and maximum values. Violin plots with smoothing in **(D)** show the median (black dotted line) and quartiles (gray lines). Statistical tests used: **(H)** Brown-Forsythe and Welch test with Dunnett’s T3 multiple comparisons correction; **(D)** Kruskal-Wallis test with Dunn’s multiple comparisons correction; **(B,C**) unpaired, two-tailed Welch’s tests; **(J)** unpaired, two-tailed Mann-Whitney test. Statistical significance is indicated as ns (p ≥ 0.05), * (p < 0.05), ** (p < 0.01), *** (p < 0.001), **** (p < 0.0001).

Since inhibiting EGFR signaling reduced microglial motility, we asked whether activating this pathway could induce directed microglial migration. To test this possibility, we injected recombinant human EGF (hEGF) into the midline of the OT at 4 dpf (**Figure 6E**, red region). This elicited robust, directed migration of microglia toward the injection site, demonstrating that hEGF serves as an attractive cue for microglia (**Figure 6F-H, Video S4**). Thus, EGFR activation is sufficient to drive microglial chemotaxis toward an ectopic hEGF source.

This finding in zebrafish prompted us to test if increasing the dose of LIN-3 EGF in the *C. elegans* germ cells is sufficient to induce their engulfment. Germline-specific overexpression of LIN-3 EGF (*Pmex-5::lin-3*) resulted in a slight increase in the number of engulfed cells in a wild-type background (**Figure S11B,C**). Surprisingly, even in apoptosis-deficient *ced-3(lf)* caspase mutants, ectopic *lin-3* expression was sufficient to induce the engulfment of a subset of non-apoptotic germ cells (**Figure 6I,J**). Notably, these engulfed cells did not show the characteristic morphology of apoptotic corpses (**Figure S11A**).

Taken together, gain-of-function experiments show that EGF ligands are sufficient to trigger efferocytosis in both models, promoting directed microglial mobility in zebrafish and apoptosis-independent engulfment in *C. elegans*. These results suggest that EGFR signaling plays a conserved role during efferocytosis in metazoans.

## Discussion

Efferocytosis is a complex, multi-step process requiring the precise regulation of diverse cellular functions. Phagocytes coordinate signals from the surroundings with their intracellular state to mount an efficient efferocytic response. This requires phagocytes to detect long-range “find-me” signals for chemotaxis and short-range “eat-me” signals for apoptotic cell recognition and uptake^10^. Phagocytes must also coordinate engulfment with degradation, balancing the uptake of new apoptotic cells with the processing of already internalized cargo^54^. The removal of apoptotic cells is therefore a complex process that may require an additional layer of regulation to coordinate the interdependencies of the individual steps and adjust overall clearance capacity. Interesting recent work has shown that macrophages can rapidly remodel their transcriptional programs via RNA polymerase II pause-release in response to corpse encounter, thereby sustaining efficient serial efferocytosis^55^.

Here, we identify EGFR signaling as such a regulatory mechanism. Rather than controlling a single step of efferocytosis, EGFR activity impacts several phases of apoptotic cell clearance. In both the *C. elegans* germline and the zebrafish optic tectum, EGFR inhibition reduces apoptotic cell recognition and uptake while also delaying the degradation of internalized corpses. Thus, EGFR signaling appears to provide a mechanism by which phagocytes coordinate efferocytosis as an integrated cellular program.

### LIN-3 EGF is an engulfment signal from germ cells that activates LET-23 EGFR in sheath cells

In *C. elegans*, *let-23 egfr* signaling controls diverse processes, such as developmental cell fate decisions^56,57^, morphogenesis^58,59^, sleep regulation^27,60^, and ovulation^29^. Our findings extend this list by identifying a new function of LET-23 modulating efferocytosis in gonadal sheath cells, in addition to its previously reported function during ovulation. The endogenous LET-23::GFP reporter is not only expressed in the sheath cells of the proximal gonads, where ovulation occurs, but also in the distal sheath cells, which mediate apoptotic germ cell clearance. Long-term imaging of *distal let-23ko* animals revealed prolonged persistence of engulfed germ cells and reduced formation of new phagosomes, pointing to a defect in apoptotic corpse recognition and degradation. A similar phenotype was observed in *somatic mpk-1(lf)* mutants. However, the reduced rate of phagosome formation in *somatic mpk-1(lf)* mutants was less pronounced, suggesting that LET-23 may interact with additional intracellular signaling pathways in engulfment. One possibility is that MAPK signaling induces a transcriptional response that modulates the phagocytic competence of the sheath cells. In addition, LET-23 may generate a localized intracellular signal, for example via the phosphoinositide signaling pathway, to produce a spatially restricted response required for apoptotic cell recognition.

Tissue-specific RNAi experiments indicated that LIN-3 EGF functions as an "eat-me" signal produced by germ cells. Interestingly, an endogenous bicistronic *lin-3* reporter displays uniform *lin-3* transcription in all germ cells. By contrast, a translational LIN-3 reporter shows a detectable signal at the cell cortex only in a subset of germ cells. This suggests that germ cells undergoing apoptosis may selectively process or secrete LIN-3 protein. Together, these results suggest that apoptotic cells may actively secrete LIN-3 towards the engulfing sheath cells, thereby contributing to their clearance in combination with other “eat-me” signals, such as PS.

Germline-specific LIN-3 overexpression in *ced-3(lf)* caspase mutants caused the engulfment of a subset of non-apoptotic germ cells. However, these germ cells accounted for only a small fraction of the total cells in the late pachytene region. One possible explanation is that a balance between pro- and anti-engulfment signals governs this process. When analyzing the AnxV::GFP reporter to detect PS exposure, we noticed a substantial heterogeneity in signal intensity among individual germ cells. Therefore, LIN-3 overexpression may promote the engulfment of germ cells that stochastically display higher PS levels. Supporting the notion of competing pro- and anti-engulfment signals, we sporadically observed engulfed, morphologically healthy germ cells in *ced-3(lf)* caspase single mutants that lack apoptosis. Thus, healthy germ cells with elevated PS exposure may exceed a threshold for activating engulfment receptors, occasionally leading to their engulfment. Overexpression of LIN-3 may further tip this balance, thus increasing the likelihood of engulfment events.

In contrast, we did not detect LET-23::GFP enrichment around somatic corpses during embryonic or larval development. Unlike the germline, non-specialized neighboring cells are responsible for clearing the individual corpses in the soma. Thus, EGFR signaling may be used only in specialized sheath cells that must operate under conditions of high apoptotic load.

### EGFR signaling promotes apoptotic cell recognition and degradation by microglia in zebrafish

In contrast to the stationary, epithelial-like sheath cells in the *C. elegans* gonads, microglia in the zebrafish OT are professional phagocytes that rely on directed cell migration to collect apoptotic neurons. Despite these fundamental differences, EGFR signaling appears to play analogous roles in apoptotic corpse recognition and clearance in both cell types.

Similar to the defects observed in *C. elegans*, pharmacological inhibition of EGFR signaling in zebrafish larvae led to the accumulation of phagolysosomes in microglia, suggesting impaired corpse degradation. Since EGFR internalization increases lysosomal protein biogenesis and lysosome number^61^, EGFR signaling may contribute to efficient corpse degradation through RAS/MAPK-dependent transcriptional upregulation of genes encoding lysosomal proteins.

In both *C. elegans* and zebrafish, the increased number of uncollected apoptotic cells after EGFR inhibition suggests an additional role for EGFR signaling in corpse recognition. Long-term imaging revealed that the engulfment rates for these two cell types are surprisingly similar. In wild-type *C. elegans*, the sheath cells engulfed an average of 6.6 apoptotic cells per gonad arm per hour, corresponding to approximately 1.6 successful engulfment events per phagocyte and hour. In comparison, zebrafish microglia engulfed an average of 2.2 apoptotic neurons per phagocyte per hour. While the *C. elegans* sheath cells are not classified as professional phagocytes, their relatively high efferocytosis rate and the fact that they are the only cells clearing a large number of apoptotic germ cells suggest that they may also be considered specialized phagocytes^1^.

Ectopic EGF also acts as an attractive cue in zebrafish, similar to LIN-3 EGF in the *C. elegans* germline. However, in zebrafish microglia, EGF promotes chemotactic migration over a relatively long distance (up to 100µm), resembling a “find-me” signal rather than an “eat-me” signal. EGF exists in both membrane-bound and soluble forms^62^. In its soluble form, EGF may form a long-range gradient guiding microglia towards apoptotic corpses, whereas the membrane-bound form could be involved in contact-dependent recognition and engulfment^63^.

### EGFR signaling orchestrates the different steps of efferocytosis

Despite the inherent differences between the *C. elegans* sheath cells and zebrafish microglia, EGFR signaling regulates the different steps of apoptotic cell clearance in a similar manner in both models. These findings are reminiscent of the functions EGFR plays in sleep regulation. EGFR signaling was first shown to regulate behavioral quiescence in *C. elegans* via a specific neuronal circuit^60^, but this process was initially considered distinct from sleep in higher animals. However, subsequent studies revealed a conserved role for EGFR signaling in sleep across metazoans ^49,60^.

EGFR signaling regulates several cellular processes necessary for efferocytosis, including cell migration, actin cytoskeletal reorganization and endocytic trafficking. This suggests several potential points at which EGFR signaling could modulate apoptotic cell clearance. Endocytosis and intracellular signaling are closely interconnected: the endocytic network regulates signaling pathways, while signaling pathways modulate endocytic trafficking^64^. In particular, EGFR endocytosis directly affects lysosomal pathways^61^.

Several observations support the idea that efferocytosis primes phagocytes and reshapes their subsequent clearance capacity through context-dependent feedback loops. In *Drosophila*, engulfment of apoptotic corpses induces a form of “molecular memory”, in which macrophages exhibit enhanced wound responses following initial corpse clearance^65^. In other cases, efferocytosis increases the expression of receptors involved in apoptotic cell recognition, such as TAM receptors^66^. These observations support a model in which the engulfment of apoptotic cells alters gene expression to modulate phagocyte function^54,67^, suggesting that phagocytes possess an adaptive toolkit that controls phagocytic capacity. Thus, phagocytes exhibit a high degree of adaptability, allowing them to dynamically adjust their responses to changing cellular and environmental contexts. In particular, inflammatory or infectious environments may dramatically alter phagocyte responses^68,69^. For example, Pan et al. (2025) reported that EGFR deficiency in the myeloid lineage in a mouse model of acute ischemic kidney disease accelerates efferocytosis^70^. However, this phenotype was observed in a highly inflammatory injury context, which differs substantially from the physiological cell death occurring in the *C. elegans* and zebrafish models. Thus, EGFR may promote efferocytosis in a homeostatic context during normal development. In contrast, in an inflammatory setting, a loss of EGFR signaling may trigger compensatory pathways that enhance phagocytic activity via inflammatory signaling.

In summary, our results demonstrate that EGFR signaling orchestrates the different steps of efferocytosis to regulate overall phagocytic capacity across metazoans. By tightly controlling localized EGFR signaling, phagocytes can adapt their ability to recognize, engulf and degrade apoptotic cells to different environmental conditions.

### Limitations

Here, we identify EGFR signaling as a conserved regulator of efferocytosis in two evolutionarily distant in vivo systems. Still, we have not yet determined whether this requirement extends to mammalian phagocytes. Testing this in mammalian cell lines may be complicated by extensive alterations in growth factor signaling in immortalized cell lines and by compensatory mechanisms that can mask the effects of EGFR pathway perturbations^71^. In addition, our data suggests that EGFR signaling is particularly important under conditions with high apoptotic load. Future work will be needed to determine whether the pathway is also required when the apoptotic cell burden is low.

## Materials and Methods

### C. elegans: culture and strains

*C. elegans* strains were maintained at 20°C on NGM plates seeded with OP50 as previously described^72^. The *C. elegans* strain N2 (Bristol) was used as wild-type reference. Before an experiment, worms were maintained in a non-starved state at 20°C for at least three generations. If not otherwise described in the figure legends, *C. elegans* hermaphrodites were synchronized at the L4 stage by picking and imaged 24 hours post-synchronization (24hpL4). For some experiments, larvae were initially synchronized at the L1 stage by washing plates containing mixed-stage worms and treating them with a bleaching solution containing 10% sodium hypochlorite (425044, Sigma-Aldrich) and 10% 5M NaOH. Worms were washed three times with M9 buffer and allowed to hatch overnight in a 15 mL Falcon tube containing 5 mL M9 buffer while shaking. L4-stage worms were then picked at 48h post-L1 for further synchronization.

For each experiment, the raw data are shown in **Data S1**. A list of all *C. elegans* strains used in this study is provided in **Data S2** (strain list). Alleles are first introduced in the text by allele number and functional class (e.g., *ced-3(n717lf)*) and subsequently referred to by functional class only (e.g., *ced-3(lf)*). For each gene, one allele per function class was used.

### C. elegans: Strain Construction

The alleles *zhIs176*, *zhIs207*, *zhIs209*, *zhIs213* and *zhIs214* were generated using the split hygromycin CRISPR-based approach^73,74^, resulting in single-copy transgene insertions at defined landing pads on chromosomes I, II or III. The endogenously tagged *lin-3(zh185)* and *let-23(zh138)* alleles were generated using the Self-Excising Drug Cassette (SEC) CRISPR approach^75^. The allele *zhEx680* was generated as an extrachromosomal array^76^. Repair template plasmids for both the split hygromycin and SEC approaches were created using Gibson assembly^77^. The sgRNA and Cas9 plasmid for the *lin-3(zh185)* endogenous tag was made using cycle-restriction ligation^78^. Details of cloning procedures, including templates and primers, are provided in **Data S3**.

Adult hermaphrodite germlines were microinjected using a DNA-based injection mix consisting of a repair template plasmid and a plasmid expressing the sgRNA under a PolIII promoter together with Cas9 under a global *eft-3* promoter. For the SEC approach, the co-injection marker plasmids pCFJ90, pCFJ104 and pGH8 were included. The extrachromosomal array *zhEx680* was generated by injection of the *Punc87b::let-23* construct together with the pCFJ90 co-injection marker. The injection mixes for all constructs are listed in **Data S3**.

### C. elegans: Microscopy

Most imaging experiments were performed on pads containing a 4% agarose dissolved in H2O (7-01P02-R, BioConcept). Worms were picked and anesthetized in 5 µl of 20 mM Tetramisole (T15-12, Sigma-Aldrich) on an agarose pad, then covered with a coverslip. *C. elegans* were imaged using three different microscopes. The microscope and objectives used for each experiment are listed in **Data S1**. (1) Most images were acquired using a Leica DMRA upright microscope (Leica, Switzerland) equipped with two sCMOS cameras (C11440-42U30, Hamamatsu, Japan), an image splitter (TwinCam, Cairn Research) for simultaneous dual-channel epifluorescence imaging and a multicolor fluorescence light source (Spectra, Luminor). Brightfield field images are acquired using a multicolor LED (B200-RGB, ScopeLED, USA). Z-stacks were acquired using a piezo objective drive (MIPOS 100 PL SG, Piezosystems Jena, Germany). Images were acquired using the HCX PL APO 40X/NA 1.32 OIL objective. (2) Long-term imaging experiments were performed using a Leica DMRA2 upright microscope (Leica, Switzerland) equipped with a sCMOS camera (Prime BSI, Photometrics, USA), a multicolor LED (Spectra, Lumencor, USA), a brightfield white LED (MCWHLP1, Thorlabs, USA), and a piezo objective drive (MIPOS 100 SG, Piezosystems Jena, Germany). Temperature control during long-term image acquisition is achieved using the custom-fitted cage incubator (H201-T-UNIT-BL_CRYO and H201-ENCLOSURE-CRYO, Okolab S.r.L., Italy). All time-lapse imaging was acquired using the HCXL PL APO 40x/NA 1.30-0.60 oil objective. (3) Confocal images were acquired using a BX61 (Olympus) microscope equipped with an X-light V2 spinning disc unit (Crest optics, Italy), an EMCCD camera (iXon Ultra 888, Andor, UK), and two high-power LEDs (UHP-T-460-DU and UHP-T-560-DI, Prizmatix, Israel) for GFP and RFP excitation. For DAPI excitation, a mercury-vapor lamp (X-Cite Exact, Excelitas Technologies Corp., USA) was used. Brightfield field images are acquired using a white LED (MCWHLP1, Thorlabs, USA). Z-stacks were acquired using a piezo objective drive (MIPOS 100 PL SG, Piezosystems Jena, Germany). Images were acquired using a 40x oil objective (Olympus UplanFL N 40x/NA 1.30), a 60x oil objective (Olympus UPlanAPO 60x/NA 1.40) or a 100x oil objective (Olympus UPlanAPO 100x/NA 1.40). All microscopes are operated using custom-built MATLAB-based imaging software (MATLAB R2023b, MathWorks, USA). Image acquisition on the microscopy systems was coordinated using a microcontroller (Arduino Mega 2560, Italy).

Images were analyzed using Fiji^79^. For visualization purposes only, *C. elegans* images were deconvolved using a custom MATLAB script that adapted the YacuDecu implementation of CUDA-based Richardson-Lucy deconvolution^80^. Synthetic point spread functions (PSFs) were generated using a Gibson-Lanni optical model^81,82^.

### C. elegans: Time-lapse Imaging

For the experiments shown in **Figures 1J** and **3G**, the lifetime of CED-1::GFP-labeled corpses was quantified in control, *distal let-23ko,* and *somatic mpk-1(lf)* strains. Time-lapse imaging was performed using microfluidic imaging chips as previously described^36,83^. The single-worm trap chambers on the chip maintained worm identity and orientation. Worms were continuously supplied with an NA22 bacterial suspension via the microfluidic device’s food inlet. Worms were loaded into the devices as young adults (after the final larval molt) and imaged overnight. Track start 0 was defined as the time of appearance of the first engulfed corpse, which in our dataset typically occurred closely in time to the first ovulation. Image stacks were acquired every 5 minutes, and the first 8 hours after the first engulfed corpse appeared were used for analysis. CED-1::GFP-positive engulfed corpses were manually tracked using the TrackMate Fiji plugin^84^. To count the number of engulfment events per hour in **Figure 3G**, the first time point where a corpse became fully engulfed by the CED-1::GFP signal was used as the start point of 6 hours of analysis.

### C. elegans: Gonad Dissection

Worms were transferred at 24 hpL4 into an indented glass dish containing 500 µL PBS supplemented with 0.2 mM tetramisole. Gonads were dissected by decapitating the worms with a syringe needle, which releases the gonads from the body cavity. After a maximum of 5 minutes in the dissection dish, dissected worms were transferred with minimal liquid into 500 µl of PBS containing 0.05% Tween (P1379, Sigma-Aldrich) and 4% Formaldehyde (F8775, Sigma-Aldrich) in siliconized 1.5 ml Eppendorf tubes. After fixation for 20-30 minutes, dissected worms were washed three times with 1 ml PBS containing 0.05% Tween (PBS-T). All washing steps were carried out in siliconized tubes, and gonads were collected by centrifugation at 800 rcf. Samples were incubated in 0.1 µg/ml DAPI solution for 10 minutes at room temperature, gently shaking in the dark. Two additional wash steps with 1 ml PBS-T were performed before mounting the dissected gonads on glass slides in 50% Mowiol solution (81381, Sigma-Aldrich) and covering them with a coverslip. Slides were incubated for 15 minutes at room temperature in the dark, then stored at 4°C until the Mowiol had polymerized.

### C. elegans: Auxin Inducible Degradation

1 mM auxin (Indole-3-acetic acid, I3750, Sigma-Aldrich)-containing NGM plates were prepared as described^35^. Auxin plates were seeded with OP50, and worms were transferred to auxin plates either for 18 h or 6 h before imaging at 24hpL4. As a control, the same *let-23::AID* strain was transferred to NGM plates containing the solvent (0.0625% EtOH) without auxin.

### C. elegans: RNA Interference

RNA Interference (RNAi) was performed by feeding worms *E. coli* HT115 bacteria producing dsRNA targeting the gene of interest. Bacteria were grown overnight at 37°C in 3 mL of 2xTY medium containing 100 µg/mL ampicillin (A6352, ITW Reagents). The following day, 1 ml of the overnight culture was mixed with 1 ml of fresh 2xTY medium containing 100 µg/ml ampicillin and 1 mM IPTG (A1008, ITW Reagents), and incubated for 3-5 hours at 37°C before seeding. NGM plates containing 1 mM IPTG and 100 µg/ml ampicillin were seeded with 300 µl of the IPTG-induced bacterial culture. *C. elegans* strains were grown on *E. coli* OP50 and transferred to RNAi plates at the L4 stage for 24 hours before imaging. Empty-vector (*ev*) bacteria containing the L4440 plasmid backbone without a gene-specific insert were used as a control. *rpn*-*6* RNAi was used as a positive control to verify RNAi efficacy for each plate batch. All RNAi clones used in this study are part of the Ahringer RNAi library^85^. For soma-specific RNAi, a strain carrying the *ppw-1(pk1425)* mutation, which results in germline-specific RNAi deficiency, was used^86^. The germline-specific RNAi strain carries a germline-specific *rde-1* rescue transgene in the *rde-1(lf)* RNAi-deficient background^87^.

### C. elegans: Scoring Germ Cell Engulfment

Engulfed germ cells were scored using the engulfment reporters *ced-1::gfp* (*bcIs39[Plim-7::ced-1::gfp]*^31,32^)*, PIP2::mCherry* (*zhIs198[Plim-7::ΔPpes-10::mCherry::PH(PLCdelta1)::unc-54 3’UTR::Loxp::Prps-0::hygromycinBres::unc-54 3’UTR::Loxp]*)^26^*, or PIP2::gfp* (*zhIs209[Plim-7::ΔPpes-10::gfp::PH(PLCdelta1)::unc-54 3’UTR::Loxp::Prps-0::hygromycinBres::unc-54 3’UTR::Loxp]*).

All experiments in which engulfment was scored were conducted at 24hpL4. One gonad per hermaphrodite was analyzed. In more than 90 % of worms, one gonad arm is positioned on top of the intestine and the other below. The germline on top of the intestine was used for analysis. Z-stacks covering the entire apoptotic zone of the germline were acquired, and engulfed cells were counted manually through the Z-stack. Cells were scored as engulfed if they were enclosed by the engulfment reporter by at least three-quarters of the cell. For larger phagosomes, DIC images were used to count the number of engulfed cells per phagosome. For **Fig. 5E,F**, cells were only counted as NUC-1::mCherry-positive if the NUC-1::mCherry signal surrounded a corpse (e.g., three pink arrows in the top right of the *let-23ko* animal) or if a corpse was filled with NUC-1::mCherry signal (e.g., the two pink arrows at the bottom of the *let-23ko* animal).

### C. elegans: Offspring Analysis

Total brood size was determined as the number of offspring produced by a single hermaphrodite over 7 days of adulthood. Individual L4 hermaphrodites were transferred to fresh NGM plates and allowed to lay eggs for 24 hours before transfer to a new plate. Transfers were repeated for 4 consecutive days. Hatched progeny were counted 24 h after the mother animal was transferred. The final plate was monitored for an additional 3 days .

### C. elegans: ERK-KTR sensor

The transgene *zhIs213[Plin-31::modERK-KTR::GFP::T2A::mCherry::his-58::unc-54 3’UTR ::Loxp::Prps-0::hygromycinBres::unc-54 3’UTR::Loxp]* was used to score MPK-1 ERK activity in the vulval precursor cells (VPCs). To image the modERK-KTR sensor in the VPCs at the L3 larval stage, L1 larvae were synchronized by bleaching and plated, then allowed to grow for 24 hours. Worms were prepared for imaging and loaded into microfluidic short-term imaging chips as described^58^. Images were flat-field corrected. To quantify the normalized activity ratio in the VPCs, the mCherry::HIS-58 nuclear reporter was used to generate nuclear masks. The average modERK-KTR::GFP intensity within the nuclear mask was divided by the average mCherry::HIS-58 intensity in the same nuclear mask. The ratio was calculated from a summed-intensity projection of the in-focus ±1 slices. The resulting ratio for each VPC was normalized to the average activity ratio for all VPCs in a given worm, whereas P3.p was excluded from the analysis, as this cell frequently undergoes fusion with the hypodermis^88^. The transgene *zhIs214[Plim-7::Δpes-10::modERK-KTR::GFP::T2A::mCherry::his-58::unc-543’UTR::Loxp ::Prps-0::hygromycinBres::unc-54 3’UTR::Loxp]* was used to score MPK-1 ERK activity in the adult sheath cell pairs 1-4. To calculate the cytoplasmic-to-nuclear ratio of the modERK-KTR::GFP construct in the sheath cells, a flat field correction was applied to the images before intensity measurements. One nucleus per sheath cell pair was measured. The ratio was calculated from a single z-slice in which the individual nucleus was in focus. The mCherry::HIS-58 nuclear reporter was used to determine nuclear masks. The average modERK-KTR::GFP intensity was measured within the nuclear mask and in the surrounding cytoplasm, as shown in **Figure S4**.

### Zebrafish: culture and drug treatment

Zebrafish (*Danio rerio*) were raised, maintained and bred according to standard procedures as described in “Zebrafish – a practical approach”^89^. All larvae used for experiments were younger than 5 days post fertilization (dpf), in accordance with the European Union Directive 2010/63/EU and local authorities (Kantonales Veterinäramt, Fishroom license TVHa Nr. 178). At the larval stages analyzed, sex determination had not yet occurred, and both sexes were included. Live embryos were kept at 28°C in E3 medium and staged according to Kimmel et al.^90^. Pigmentation was prevented by adding 0.003% N-phenylthiourea (PTU, P7629, Sigma-Aldrich) to E3 medium from 1 dpf until the end of the experiment. Larvae were anesthetized using 0.01% tricaine (A5040, Sigma-Aldrich). A list of all zebrafish lines used in this study is provided in **Data S3**. Zebrafish were treated with either DMSO-containing E3 medium (D2438, Sigma-Aldrich) or E3 medium containing DMSO and 20 µM Erlotinib Hydrochloride (S1023, Selleckchem), resulting in a final concentration of 1% DMSO in all conditions. Treatment initiated at 2 dpf, 24 hours before imaging. During staining and imaging, larvae were maintained in the same medium. To ensure consistent drug solubility, small aliquots of DMSO and Erlotinib stock solutions were used. Erlotinib stock solutions (5 mM in 100% DMSO) were stored at -80 °C and used within 6 months.

### Zebrafish: Acridine Orange Staining

Zebrafish larvae were stained for 1 hour in the dark at 28 °C in 10 ml E3 containing 10 µg/ml Acridine Orange (AO, A6014-10G, Sigma-Aldrich). After staining, fish were washed 6 times in 40 mL of fresh E3 medium to remove excess AO. Fish were mounted immediately after washing and imaged between 45 minutes and 2 hours after washing (corresponding to 45 minutes to 2 hours of destaining).

### Zebrafish: Microscopy

Before mounting, larvae were anesthetized using 0.01% tricaine. Fish were mounted in 1.3% low-melting agarose (PeqGOLD Low Melt Agarose, PeqLab Biotechnologie GmbH) in E3 containing 0.01% tricaine. Zebrafish larvae were mounted in glass-bottom dishes, and after the agarose had set, 1 ml of medium containing the indicated condition medium was added. All zebrafish experiments were imaged using the Andor Dragonfly 200 Sona spinning-disc microscope (Oxford Instruments, UK) with the Nikon 20x/NA 0.95 water or 40x/NA 1.25 silicon oil objectives, except the EGF injection experiment (Figure 6A-D), which were performed on the Olympus BX61 microscope (see *C. elegans* Microscopy, microscope (3)) equipped with an Olympus UPlanFI 20x/NA 0.5 air objective. Images were analyzed using Imaris (Oxford Instruments, UK) or Fiji.

### Zebrafish: Image Analysis

For analysis of uncollected apoptotic neurons in *fms::mCherry* and *fms::mCherry; irf8-/-* larvae, animals were stained with Acridine Orange (AO) as described above. 3D image volumes of the zebrafish brain were used to detect AO-positive cells using the Imaris Spot detection tool in the AO image channel. A spot diameter of 4 µm was used with quality thresholds of either 20 or 50, as specified in the figure legend. Spots detected outside of the region of interest (OT or HB) were manually removed. For *fms::mCherry* larvae, detected spots within microglia were removed manually when quantifying uncollected apoptotic cell numbers. Microglia were segmented for AO sum intensity and microglial volume quantification using the Imaris Surface tool, with machine-learning-based classification applied to the *fms::mCherry* channel. For sum intensity measurements, less stringent masks were generated to ensure inclusion of AO material within branches and peripheral phagosomes. For volume measurements, a separate model was trained using more stringent mask definitions to improve accuracy. Incomplete microglial masks due to inaccurate detection or partial imaging of the microglia not fully contained within the imaging stack were excluded and not manually corrected. Sum intensity and volume measurements per microglia were extracted using the built-in Imaris analysis tools.

Lamp1-mGFP-positive vesicles were counted and measured in 3 dpf larvae using Fiji. Each microglial cell was analyzed by traversing the stack and measuring the area and diameter of Lamp1-Positive vesicles. Only vesicles >1 µm diameter and fully surrounded by Lamp1-mGFP were scored.

### Zebrafish: time-lapse imaging

The *fms:lyn-miRFP; fms:Lamp1-mGFP* larvae were imaged for between 1 and 1.5 hours using 30-second image acquisition intervals.

For the number of successful engulfment events, only events resulting in a fully formed phagosome in the lyn-miRFP670 channel were counted. Only microglia that were within the imaging z-stack for 60 minutes were analyzed. Lamp1-mGFP-positive vesicles were tracked in 3 dpf larvae using the Fiji plugin Trackmate. The vesicles were tracked for at least 10 minutes. Vesicle sizes were measured at the start and end of tracking, and the shrinkage rate was calculated as the average change in size over 10 minutes.

### Zebrafish: EGF injections

Zebrafish carrying the *mpeg1:gfp-caax* reporter were injected at the optic tectum midline at 4 dpf. To minimize the time between injections and imaging and to facilitate easy access to the injection tissue, injections and imaging were performed on an Olympus BX61 upright microscope, allowing access to the midline from above. Larvae were mounted in custom parafilm wells prepared by melting 4 layers of parafilm with punched circular openings onto glass slides, which allowed mounting of zebrafish and addition of E3 medium onto the glass slide. Before mounting, larvae were anesthetized using 0.01% tricaine. Fish were mounted on the parafilm glass slide in 1.3% low-melting agarose prepared in E3 medium containing 0.01% tricaine. After the agarose had solidified, the agarose adjacent to the optic tectum was carefully removed with a scalpel to improve accessibility for injection. The parafilm well was filled with E3 medium containing 0.01% Tricaine. Zebrafish were injected using 20 µm diameter pre-cut injection pipettes (VZIP-OD-TL-PL, BioMedical Instruments, Germany) mounted on a micromanipulator attached to the PV820 Pneumatic Picopump (World Precision Instruments, USA). This setup allowed immediate imaging following injection. Larvae were injected with HBSS (H6648, Sigma-Aldrich) containing 0.05% phenol red (P0290, Sigma-Aldrich) as a control solution or HBSS supplemented with 0.05% phenol red and 500 nM hEGF (E9644, Sigma-Aldrich) for the EGF condition. The larvae were imaged for 30 min at 30 sec intervals following injections. Images were aligned by the midline before analysis. Microglia cell body positions in XY were tracked for 30 min after injections using TrackMate in Fiji.

### Zebrafish: Microglial Mobility

To quantify microglia mobility in larvae at 3 dpf, time-lapse z-stacks of individual microglia were manually cropped from movies to facilitate feature extraction. The smaller 4D image stacks reduced memory requirements. To ensure reliable segmentation of microglial branches and finer substructures, the *fms:lyn-miRFP670* reporter line, which shows strong membrane labeling, was used. Microglia were imaged for at least 1 hour, and z-stacks were acquired every 30 seconds. Masks were generated using a trained neural network model and subsequently skeletonized to represent the microglial branch structure. A soma mask was defined to exclude skeleton components within the microglial cell soma.

Microglial segmentation, branch dynamics and structural analysis were performed using a multi-step image analysis workflow comprising several custom scripts, as described in **Data S4** and illustrated in **Figure S10**. Code files were named according to the corresponding panel in **Figure S10**, which shows the different image processing steps.

### Statistical analysis

Statistical analyses were performed using GraphPad Prism version 10.6.1 software (GraphPad Software Inc, USA) as indicated in the figure legends. For most experiments, comparisons between two groups were performed using unpaired, two-tailed Welch’s t-tests assuming normal distribution with unequal variances. For comparisons involving three or more groups, the Brown-Forsythe and Welch ANOVA tests, followed by Dunnett T3 multiple comparisons, were used, assuming normality with unequal variances. For comparisons of two groups, which include conditions with non-normal distributions (e.g., *ced-3(lf)* mutants or offspring analysis including sterile animals), an unpaired, two-tailed Mann-Whitney test was used. For non-normal distributions with more than three groups, a Kruskal-Wallis test followed by Dunn’s multiple comparisons was used. Statistical significance is indicated as ns (p ≥ 0.05), * (p < 0.05), ** (p < 0.01), *** (p < 0.001), **** (p < 0.0001). All raw data used for statistical analyses are provided in **Data S1**.

## Supporting Information

**Supplementary Data.** Contains supplementary Figures 1-11:

**Figure S1.** Related to Figure 1 **Figure S2.** Related to Figure 2 **Figure S3.** Related to Figure 2 **Figure S4.** Related to Figure 2 **Figure S5.** Related to Figure 3 **Figure S6.** Related to Figure 4 **Figure S7.** Related to Figure 5

**Figure S8.** Related to Figure 6 **Figure S9.** Related to Figure 6 **Figure S10.** Related to Figure 6 **Figure S11.** Related to Figure 6

**Video S1.** Phagolysosome tracking in *fms:Lamp1-mGFP* reporter

**Video S2.** EGF injections in the zebrafish midline

**Video S3.** Microglial mobility in Erlotinib-treated zebrafish

**Video S4.** Animation of the microglial mobility pipeline shown in Figure S10

**Data S1**. Raw data used to generate graphs and additional information on experimental procedures

**Data S2**. Strain list for *C. elegans* and zebrafish

**Data S3.** Details on strain construction and plasmids

**Data S4.** Code documentation and scripts used in image processing for microglial mobility measurements

## Acknowledgements

We thank all past and present members of the Hajnal group for the insightful discussions and continuous support. In particular, we are grateful to Tea Kohlbrenner and Ana Laranjeira for establishing the foundation for studying germ cell apoptosis in our laboratory. We further thank Julian Bamert and Benjamin Egli for generating the PIP2::GFP construct. We thank the Peri and Gilmour groups for helpful discussions and support with zebrafish experiments. We are grateful to Cornelia Henkel for fish care and to Nathalie Tichy and Ayush Adity Pal for sharing their zebrafish lines and for additional zebrafish support. We thank Scott Wilcockson and Caroline Hill for providing the modERK-KTR plasmids and for valuable advice on their constructs. We also thank Peter Askjaer for valuable discussions and insights on tissue-specific knock-out approaches using their FLP toolbox. We acknowledge the *Caenorhabditis* Genetics Center (CGC) for providing *C. elegans* strains, which is funded by NIH Office of Research Infrastructure Programs (P40 OD010440). We are grateful for the openness and support of the *C. elegans* community, which provides an essential backbone for this work through numerous helpful discussions at the annual worm meetings. We thank the team of the Center for Microscopy and Image Analysis at the University of Zurich for their support with image analysis. This research was supported in part by the Swiss National Science Foundation (SNSF) grant no. 184792 awarded to A.H. and the Kanton of Zurich.

## Author Contributions

S.S., A.H., and L.F.C. conceived the project. L.F.C., A.H., and S.S. designed the *C. elegans* experiments. F.P., A.V., and L.F.C. designed the zebrafish experiments. L.F.C. performed the experiments, generated strains, and analyzed the data. L.F.C. developed the image analysis code with support from S.B. and A.H., S.B. developed the flatfield correction and deconvolution tool. S.B. performed and analyzed the VPC-specific modERK-KTR experiment. S.S. performed and analyzed the *Pmex-5::lin-3* experiment. M.D generated the modKTR-ERK construct, and S.S. generated the *let-23::AID* construct. A.H. provided funding and resources. L.F.C. and A.H. wrote the manuscript with support of F.P. and input from all authors.

