## Supplementary Figures 1-11 for "Conserved role of EGFR signaling in apoptotic cell recognition and processing across different phagocytes"

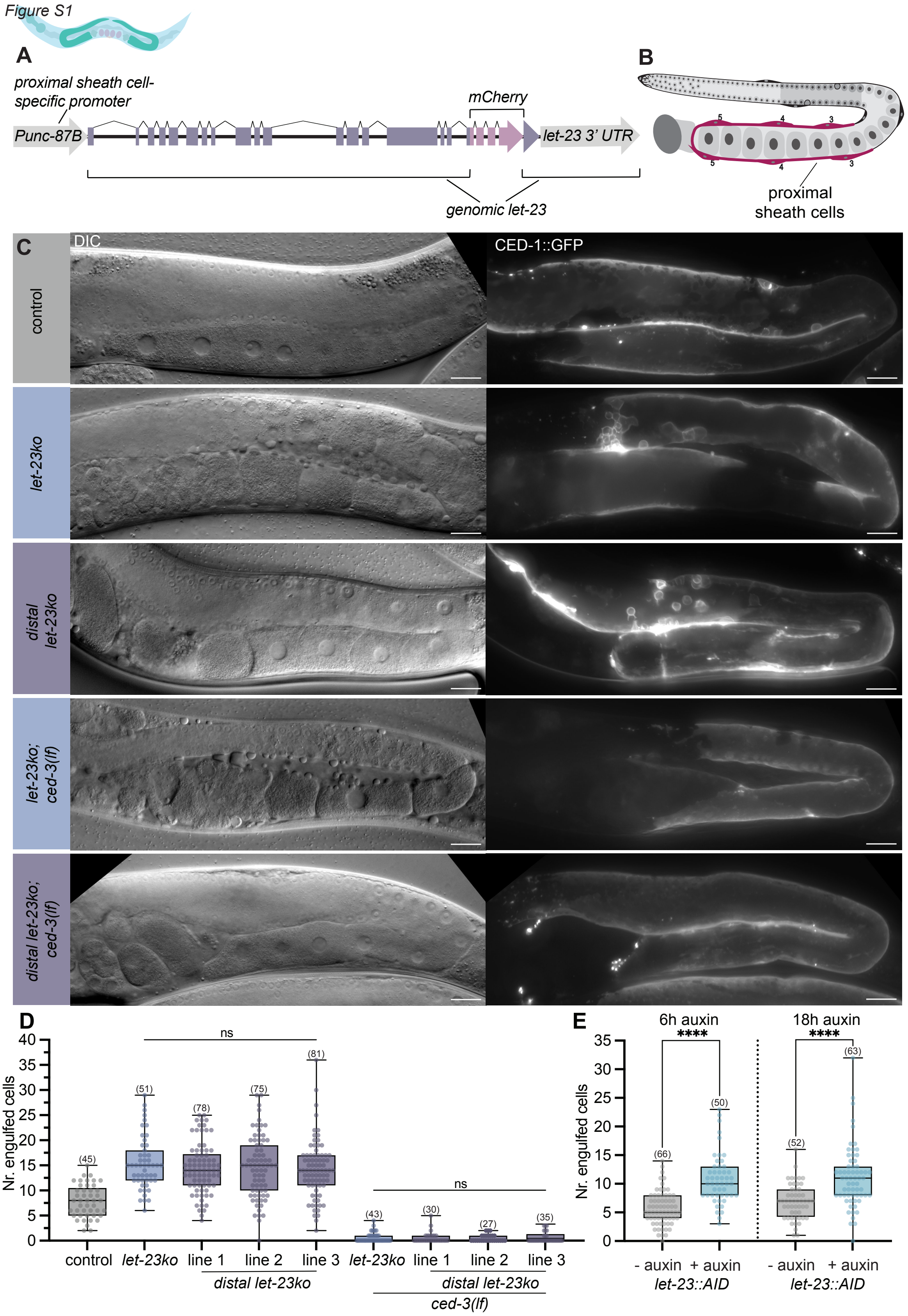

**Figure S1. Distal sheath cell-specific *let-23ko* rescue and full gonad images in *let-23ko* mutants.**

**(A)** Schematic of the proximal sheath cell-specific *Punc-87b::let-23::mCherry* construct used to rescue ovulation defects in the *let-23ko* strain (*distal let-23ko*). **(B)** Schematic of the *C. elegans* germline showing the three proximal sheath cell pairs in red. **(C)** DIC and CED-1::GFP images of the entire pachytene region and the proximal gonad in control, *let-23ko*, distal *let-23ko*, *let-23ko; ced-3(lf)* and distal *let-23ko; ced-3(lf)* animals. A single oocyte stack is visible in the DIC images of control, distal *let-23ko*, and distal *let-23ko; ced-3(lf)* animals*.* Broken-down oocytes, as a consequence of the ovulation defect, are visible in *let-23ko* and *let-23ko; ced-3(lf)* animals. Engulfed germ cells are detected by CED-1::GFP staining in maximum intensity projections. **(D)** Quantification of CED-1::GFP-positive engulfed germ cell corpses in *distal let-23ko* animals. Three independent extrachromosomal lines were scored. Differences in engulfed cell numbers between the three lines are statistically not significant. **(E)** Quantification of CED-1::GFP-positive engulfed germ cell corpses in *let-23::AID* animals (*let-23::AID*) after 6 h or 18 h of auxin-treatment. All worms shown were staged as 24hpL4. Boxplots represent the median and interquartile range, whiskers indicate minimum and maximum values. Statistical tests used: **(D)** Kruskal-Wallis with Dunn's multiple comparison correction; **(E)** Brown-Forsythe and Welch tests with Dunnett's T3 multiple comparisons correction. Statistical significance is indicated as ns ($p \geq 0.05$), * ($p < 0.05$), ** ($p < 0.01$), *** ($p < 0.001$), **** ($p < 0.0001$). Scale bars: 20 µm.

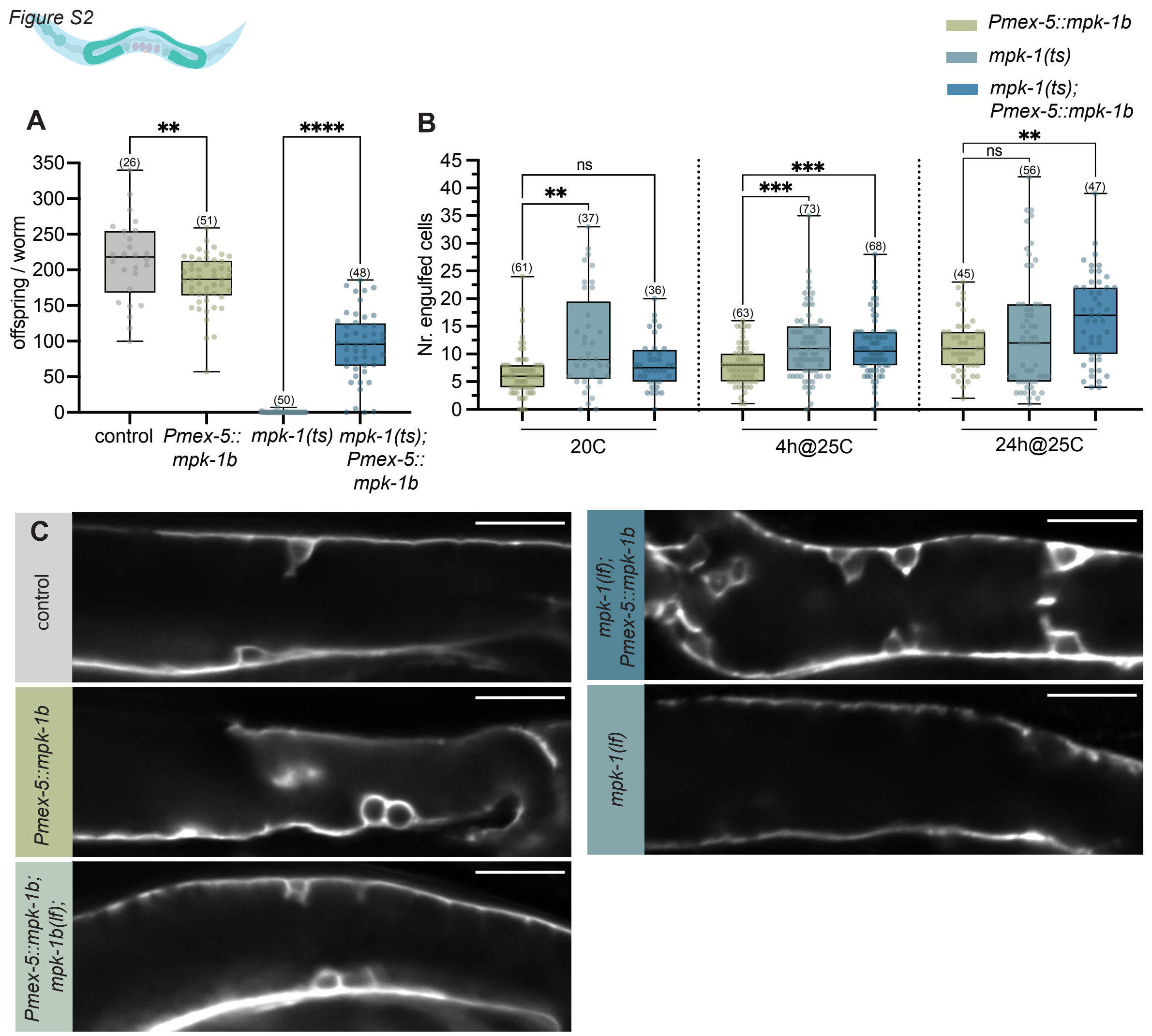

**Figure S2. Additional data on *mpk-1(ga111ts)* and *mpk-1(ga117lf)* alleles.**

**(A)** Brood size analysis of *mpk-1(ts)* temperature-sensitive mutant with germline-specific *mpk-1b* rescue (*Pmex-5::mpk-1b*). Worms were grown at the restrictive temperature of 25°C from the L1 stage onwards. Worms were transferred every 24 h to a new plate, and offspring were counted 24 h after each transfer. **(B)** Number of CED-1::GFP engulfed germ cells in indicated strains at different temperatures. All worms were analyzed 24hpL4 after varying durations at the restrictive temperature (25°C). **(C)** CED-1::GFP expression in the apoptotic zone of the indicated strains encompassing all conditions quantified in **Figure 2F**. Single z-slices are shown. All worms in this figure were staged at 24hpL4, except for **(A)**, where offspring were counted during 4 days of adulthood. The numbers in brackets in each graph refer to the number of animals scored. Boxplots show the median and interquartile range, whiskers indicate minimum and maximum values. Statistical tests used: **(A,B)** Brown-Forsythe and Welch test with Dunnett's T3 multiple comparisons correction. Statistical significance is indicated as ns ($p \geq 0.05$), * ($p < 0.05$), ** ($p < 0.01$), *** ($p < 0.001$), **** ($p < 0.0001$). Scale bars: 20 μm.

*Figure S3*

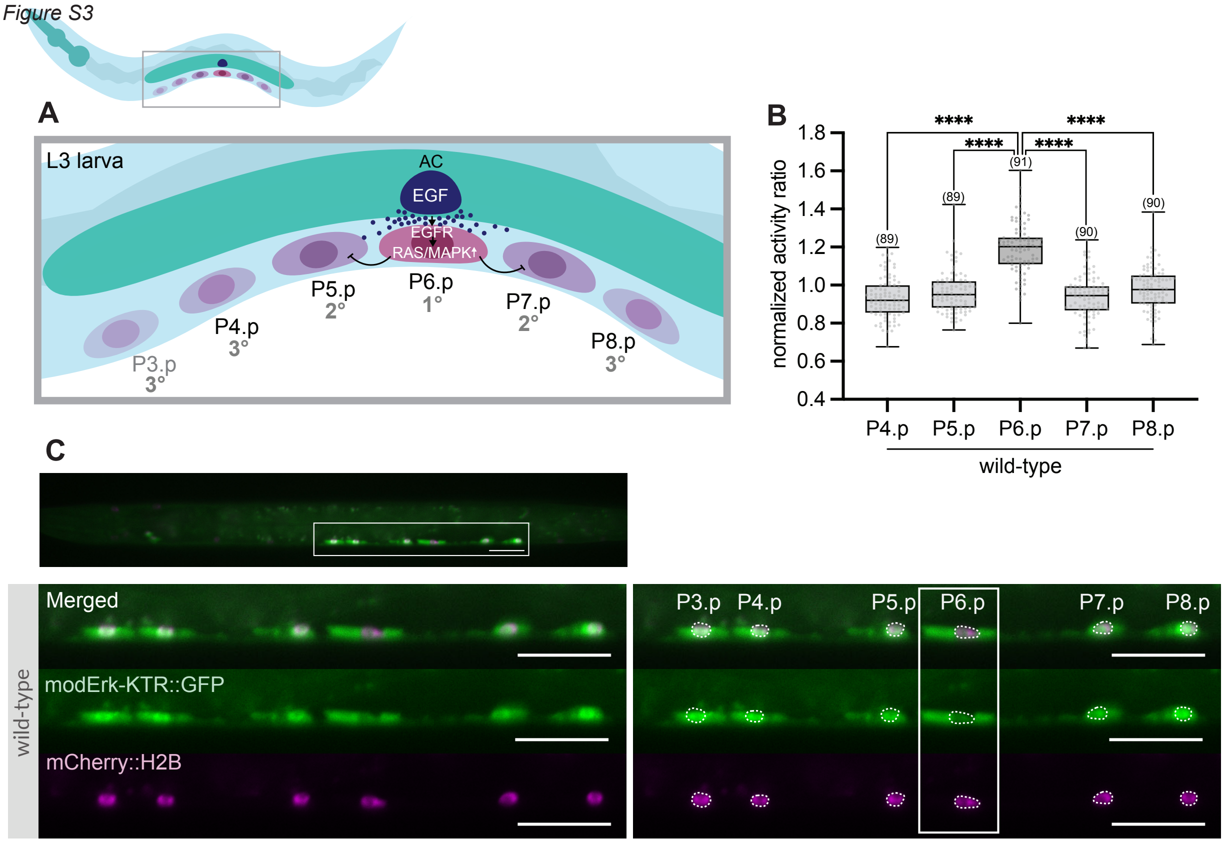

**Figure *S3*. Analysis of the modERK-KTR sensor in *C. elegans* vulval precursor cells**

**(A)** Schematic depiction of vulval precursor cells (VPCs) during vulval induction. The Anchor Cell (AC) secretes LIN-3 EGF activating the RAS/MAPK signaling cascade in the VPCs. The VPC closest to the AC, P6.p, exhibits the highest ERK MPK-1 activity. **(B)** Normalized modERK-KTR activity ratios in the VPCs. The activity ratio for each VPC except P3.p was calculated using the mCherry::H2B nuclear reporter to generate a nuclear mask. Average modERK-KTR::GFP intensity within the nuclear mask was divided by average mCherry::H2B intensity in the same region. The resulting ratio was normalized to average ratios of the 5 VPCs in each animal. **(C)** Fluorescent images of the 6 VPCs at the mid-L2 larval stage. Bottom, left: close-ups without nuclear masks. Bottom, right: with nuclear masks and cell identities. White box outlines P6.p, which exhibits the highest MPK-1 activity. All worms in this figure are staged as 24hpL1. Numbers in brackets in **(B)** refer to number of VPCs scored. Boxplots represent the median and interquartile range, whiskers indicate the minimum and maximum values. Statistical test used in **(B)**: Brown-Forsythe and Welch test with Dunnett's T3 multiple comparisons correction. Statistical significance is indicated as ns ($p \geq 0.05$), * ($p < 0.05$), ** ($p < 0.01$), *** ($p < 0.001$), **** ($p < 0.0001$). Scale bars: 20 µm.

*Figure S4*

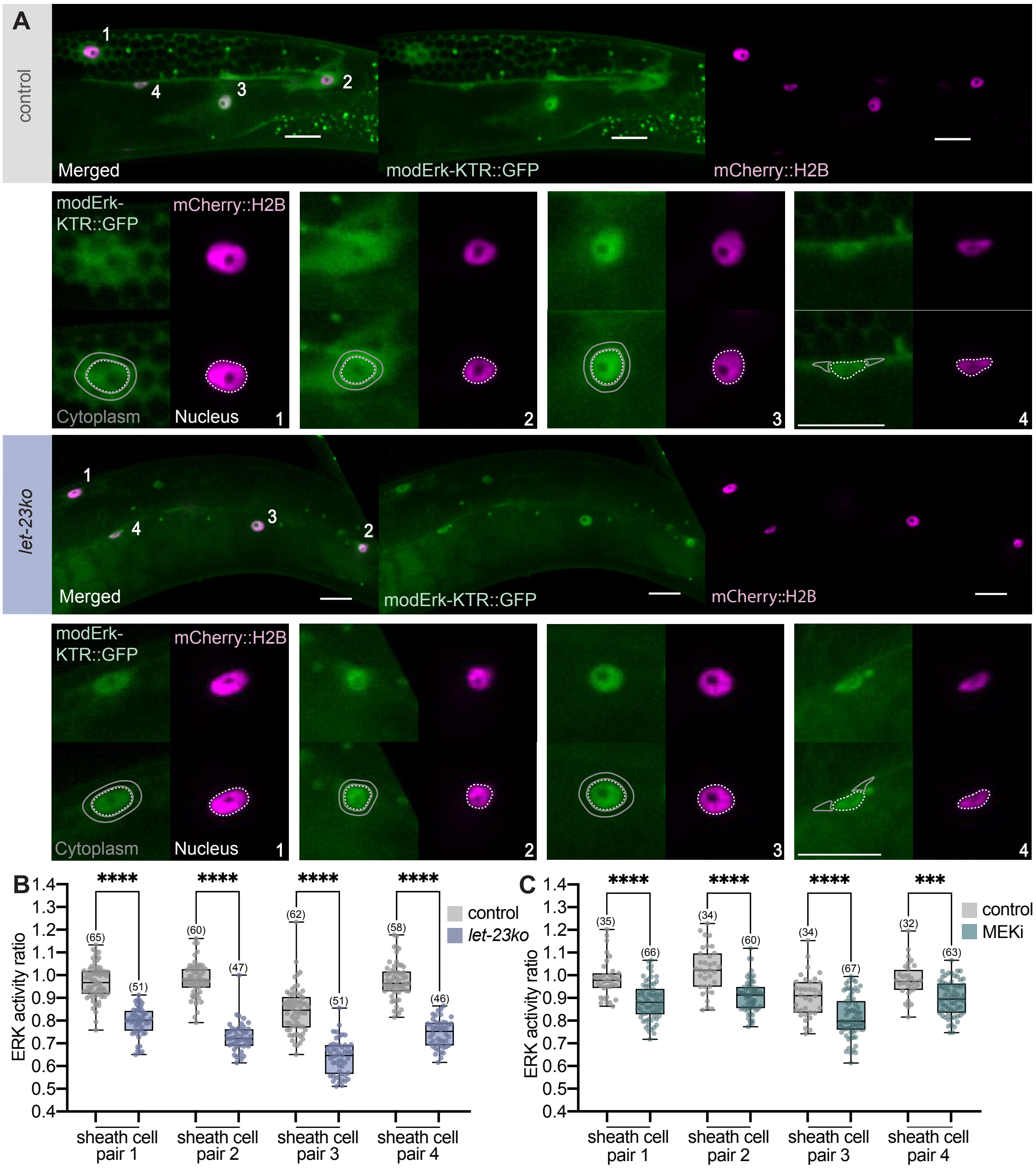

**Figure S4. Analysis of the modERK-KTR sensor in the sheath cells and MEKi drug treatment**

**(A)** Top rows: modERK-KTR (green) and mCherry::H2B (magenta) under the *lim-7* promoter in sheath cells 1 to 4 (one of each pair is shown) in wild-type control and *let-23ko* animals. Bottom rows: Close-ups of the four sheath cells expressing the modERK-KTR biosensor with the masks used for analysis. Dashed white lines show nuclear masks drawn around the mCherry::H2B signal. The solid gray lines indicate cytoplasmic masks drawn around the nuclear masks. The nuclear mask was excluded from these regions to measure cytoplasmic sensor intensity. If a nucleus was positioned laterally (e.g., sheath cell 4), the cytoplasmic signal was measured in two independent regions on both sides of the nucleus, and mean intensities of the two measurements were averaged. **(B)** Quantification of modERK-KTR biosensor in all four sheath cells in control and *let-23ko* animals, as described in methods. **(C)** Quantification of modERK-KTR biosensor activity in all 4 sheath cells in DMSO-treated control animals and after 4 h treatment with 50 µM of the MEK inhibitor U0126 in DMSO (MEKi). All worms were staged at 24hpL4. Boxplots represent the median and interquartile range, whiskers indicate minimum and maximum values. Numbers in brackets refer to the number of animals scored. Statistical tests used in **(B,C)**: Brown-Forsythe and Welch test with Dunnett's T3 multiple comparisons correction. Statistical significance is indicated as ns ($p \geq 0.05$), * ($p < 0.05$), ** ($p < 0.01$), *** ($p < 0.001$), **** ($p < 0.0001$). Scale bars: 20 µm.

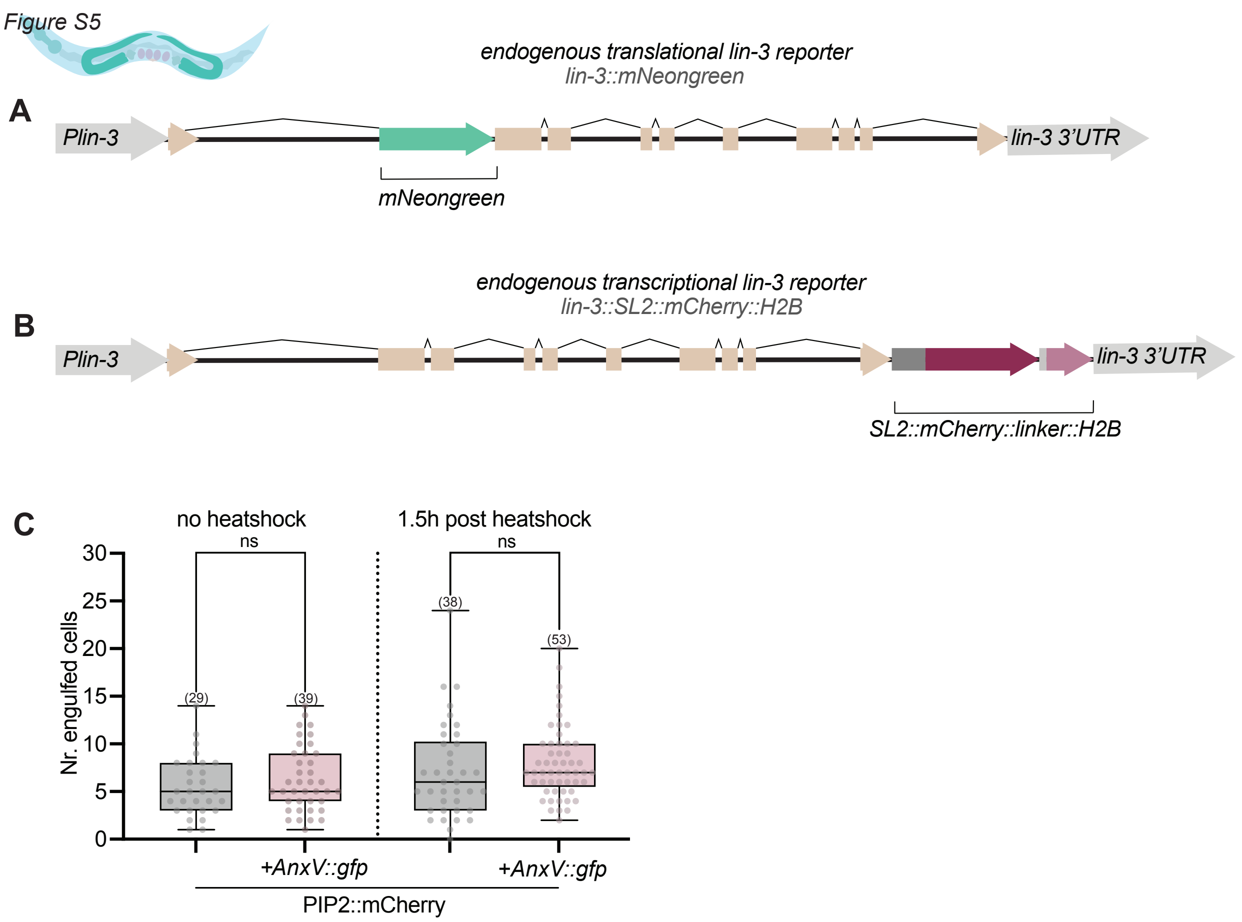

**Figure S5. Structure of the endogenous *lin-3* reporters and AnnexinV reporter control experiment.**

**(A)** Schematic of the endogenous *lin-3::mNeongreen* translational reporter[43]. The *mNeongreen* cassette was inserted before the second exon. **(B)** Schematic of the endogenous *lin-3::SL2::mCherry::H2B* transcriptional reporter. An *SL2::mCherry::H2B* cassette was inserted after the last exon. **(C)** Quantification of the total number of engulfed germ cells in worms carrying the AnxV::GFP reporter. Left: without a heat shock to induce the AnxV::GFP expression, Right: with the same heat shock conditions as used in **Figure 3F,G**. Boxplots represent the median and interquartile range, whiskers indicate minimum and maximum values. The numbers in brackets refer to the number of animals scored. Statistical test used: Brown-Forsythe and Welch test with Dunnett's T3 multiple comparisons correction. Statistical significance is indicated as ns ($p \geq 0.05$), * ($p < 0.05$), ** ($p < 0.01$), *** ($p < 0.001$), **** ($p < 0.0001$).

*Figure S6*

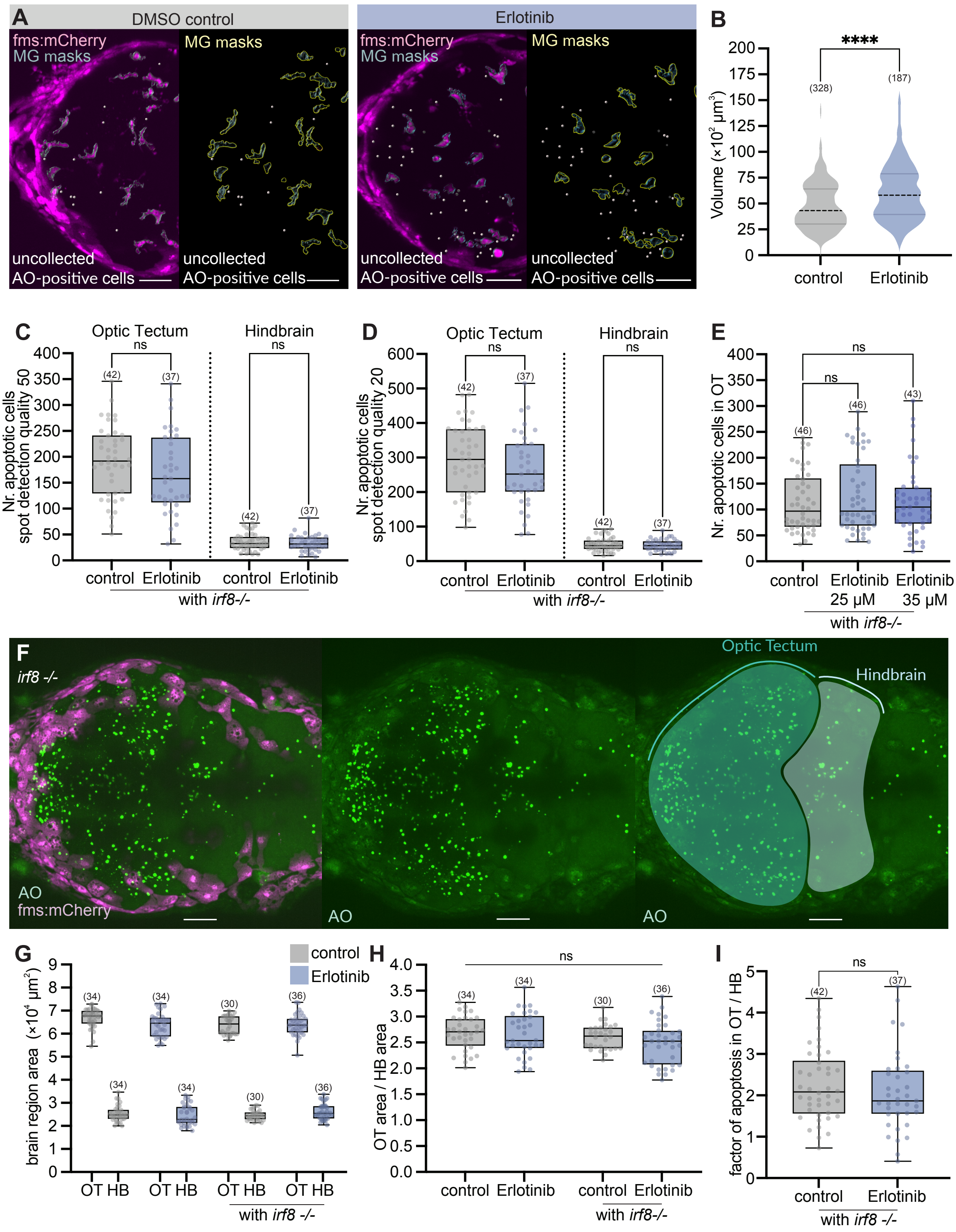
A
DMSO control
Erlotinib
fms:mCherry
MG masks
MG masks
uncollected
AO-positive cells
B
Volume (×10² µm³)
****
(328)
(187)
control
Erlotinib
C
Optic Tectum
Hindbrain
Nr. apoptotic cells spot detection quality 50
ns
control
Erlotinib
with irf8-/-
D
Nr. apoptotic cells spot detection quality 20
E
Nr. apoptotic cells in OT
Erlotinib 25 µM
Erlotinib 35 µM
F
irf8 -/-
AO
fms:mCherry
Optic Tectum
Hindbrain
G
brain region area (×10⁴ µm²)
OT HB
with irf8 -/-
H
OT area / HB area
I
factor of apoptosis in OT / HB

86

**Figure S6. Different spot detection parameters used for scoring apoptosis and brain size measurements.**

**(A)** Left: *fms:mCherry* (magenta), uncollected AO-positive apoptotic neurons (white dots) and the microglia (MG) masks (gray) in the OT of control and Erlotinib-treated zebrafish, used to measure AO staining intensity within microglia as shown in **Figure 4C**. The *fms:mCherry* reporter labels microglia in the brain and xanthophores in the skin. Right: Spot detection made in the AO channel (4 μm spot size, quality score 50), used to count uncollected apoptotic cells outside the microglia mask (yellow). See methods for details on the AO spot-detection parameters and microglia masking. **(B)** Volume per microglia in the OT in control and Erlotinib-treated zebrafish. 23 control and 27 Erlotinib-treated larvae were analyzed. **(C)** Quantification of apoptotic cells (detected in the AO channel, 4 μm spot size, quality score 50) in the OT and HB of *irf8*$_{st95}$ (*irf8-/-*) mutants treated with 20 μM Erlotinib. **(D)** Same quantification as in **(C)**, but using a lower spot detection quality threshold. **(E)** Quantification of apoptotic cells (detected in the AO channel, 4 μm spot size, quality score 20) in in the OT of *irf8-/-* mutants treated with 25 μM and 35 μM Erlotinib. **(F)** *fms:mCherry* expression (magenta) and AO staining (green) in the OT and HB of an *irf8-/-* control fish. Left: merged AO and *fms:mCherry* channels. Middle: The AO staining was used to draw masks of the OT and HB. Right: Masked OT and HB used to measure their areas. **(G)** Areas of the OT and HB in control and 20 μM Erlotinib-treated animals, in a wild-type and *irf8-/-* background. **(H)** Ratios of OT to HB area in control and 20 μM Erlotinib-treated animals, in a wild-type and *irf8-/-* background. **(I)** Apoptosis factor (apoptotic cells per area in OT divided by apoptotic cells per area in HB) in *irf8-/-* animals. The number of AO-positive cells was normalized by the average area factor of OT and HB. All zebrafish were staged as 3 dpf. All images shown are maximum-intensity projections, with the top z-slices containing skin xanthophores removed for improved visualization. Numbers in brackets refer to the number of animals scored, except for **(B)**, where the number of microglia analyzed is indicated. Boxplots represent the median and interquartile range, whiskers indicate the minimum and maximum values. Violin plot with smoothing in **(B)** shows the median (black dashed line) and quartiles (gray lines). Statistical tests used: **(B,I)** unpaired, two-tailed Welch's tests, **(C-E,G,H)** Brown-Forsythe and Welch test with Dunnett's T3 multiple comparisons correction. Statistical significance is indicated as ns ($p \geq 0.05$), * ($p < 0.05$), ** ($p < 0.01$), *** ($p < 0.001$), **** ($p < 0.0001$). Scale bars: 50 μm.

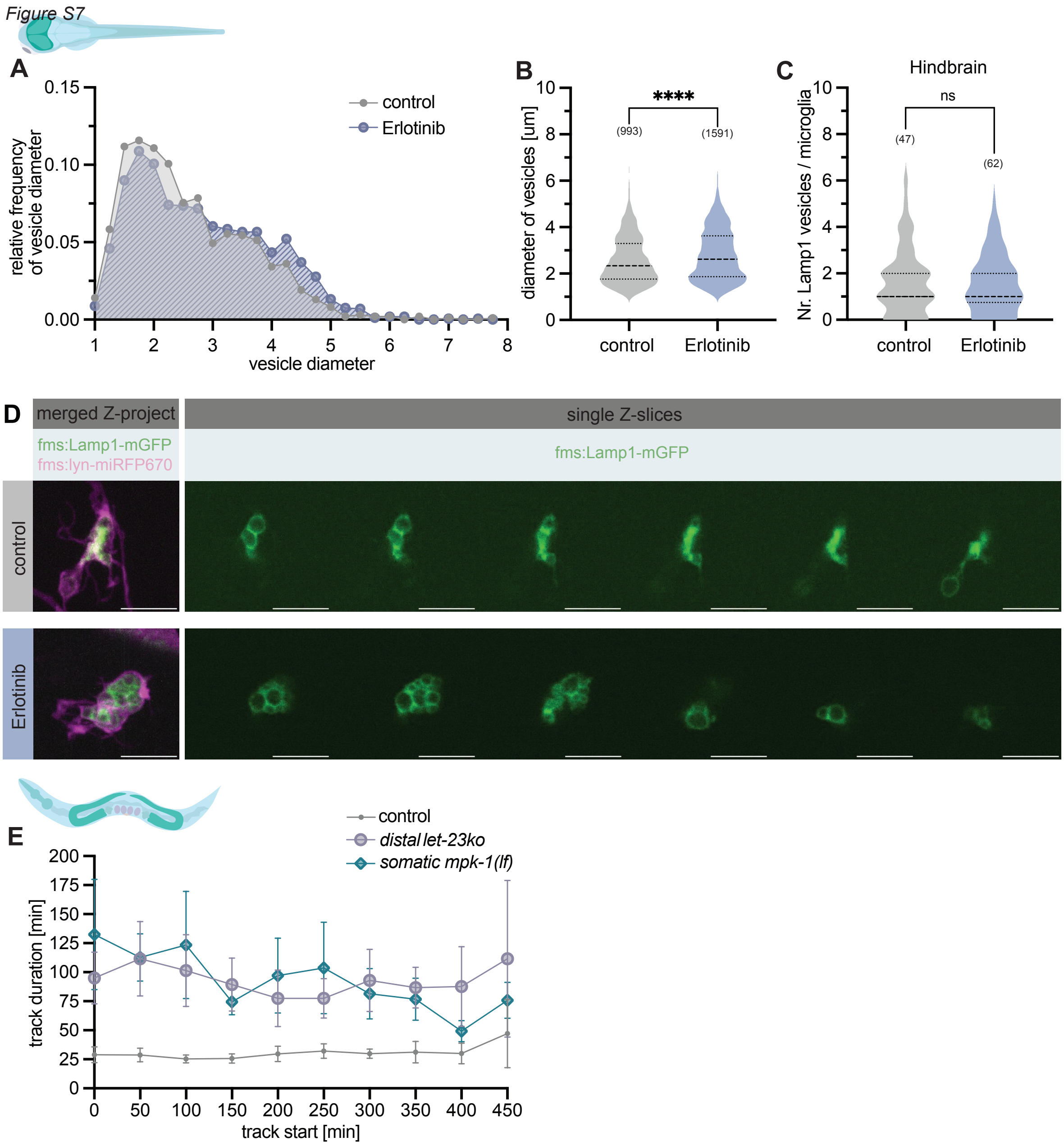

**Figure S7. Size distribution of Lamp1-positive vesicles in Erlotinib-treated zebrafish and germ cell track duration in *C. elegans*.**

**Zebrafish: (A)** Frequency distribution of Lamp1 vesicle diameters in control and Erlotinib-treated zebrafish, shown as percentage relative of total vesicles per condition. Vesicles <1 µm in diameter were excluded. Bin center steps: 0.25µm. 284 cells were analyzed in 30 control and 248 cells in 30 Erlotinib-treated animals. **(B)** Quantification of vesicle diameters in the OT of 30 control and Erlotinib-treated fish each. **(C)** Quantification of Lamp1 vesicles number per microglia in the HB of 30 control and Erlotinib-treated animals each. **(D)** Left: maximum Intensity projection of *fms:lyn-miRFP670* (magenta) and *fms:Lamp1-mGFP* (green) expression in microglia of a control and an Erlotinib-treated animal. Right:

Six individual z-slices each of *fms:Lamp1-mGFP* expression in the same cells shown to the left. All zebrafish were staged at 3 dpf.

***C. elegans:*** **(E)** Quantification of CED-1::GFP track duration relative to track start in control, *distal let-23ko* and *somatic mpk-1(lf)* worms. Six animals were recorded for each strain. Track duration is plotted against the time point at which a cell initially became CED-1::GFP-positive (track start). Timepoint 0 is defined as the first occurrence of a corpse in a recording. Numbers in brackets in **(B)** refer to the number of vesicles measured, and in **(C)** to the number of microglia analyzed. Violin plots with smoothing show the median (black dotted line) and quartiles (gray lines). Error bars in **(E)** show the standard deviation of the mean track duration. Statistical tests used in **(B,C)**: Unpaired, two-tailed Welch's tests. Statistical significance is indicated as ns ($p \geq 0.05$), * ($p < 0.05$), ** ($p < 0.01$), *** ($p < 0.001$), **** ($p < 0.0001$). Scale bars: 20 µm.

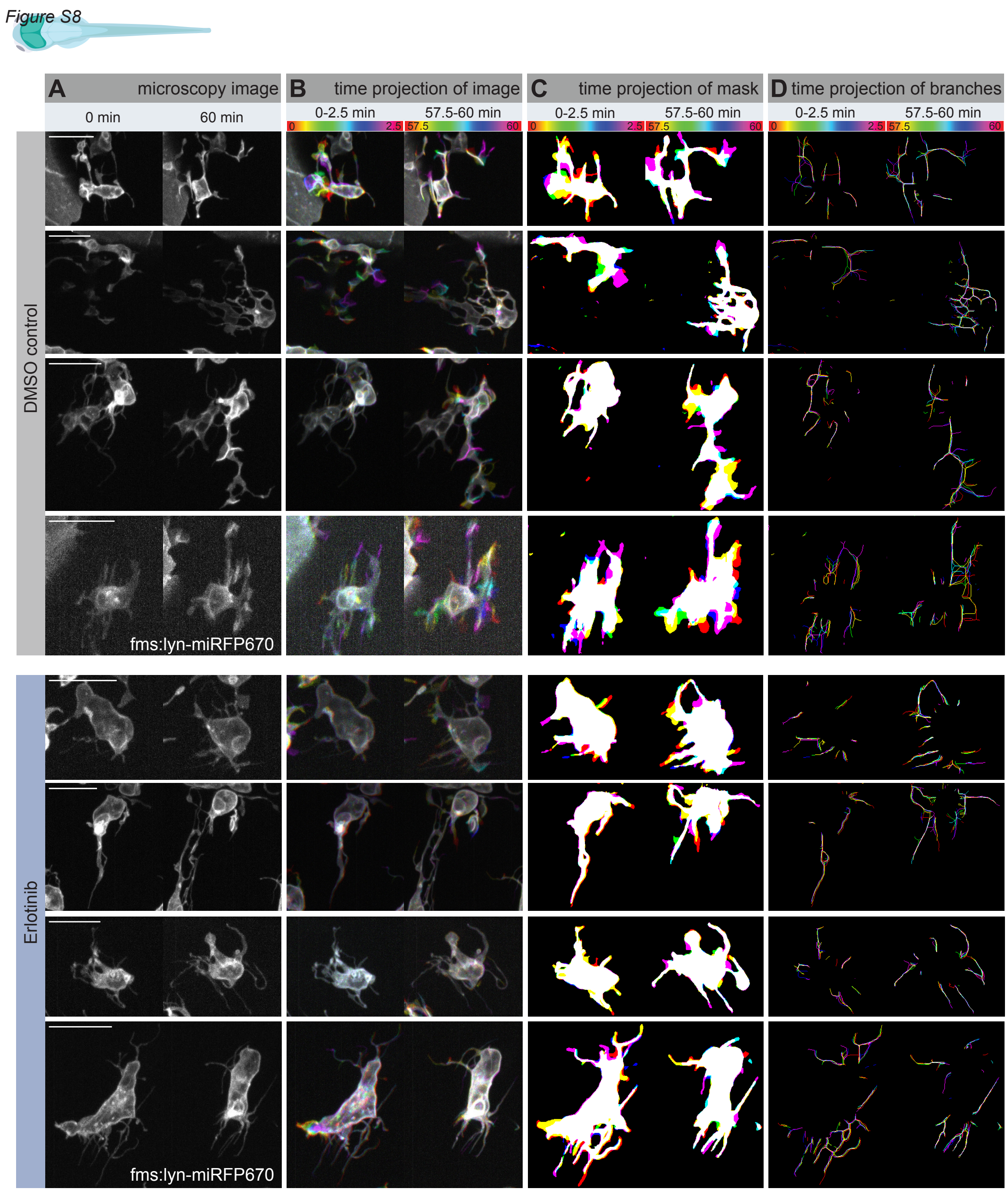

**Figure S8. Visualization of microglial mobility and branching.**

**(A-D)** Four examples of microglia in control and Erlotinib-treated 3 dpf larvae that were used for microglial mobility analysis in **Figure 6F-H** and **Figure S9**. The first example of each condition is also shown in **Figure 6E**. **(A)** shows maximum intensity projections of *fms:lyn-miRFP670* signal at the beginning and end of the 60 min recording, **(B)** color-coded time projections of five consecutive frames at the beginning and end of recording, **(C)** color-coded time projections of microglial masks, and **(D)** color-coded time projection of microglial branches. Scale bars: 20 µm.

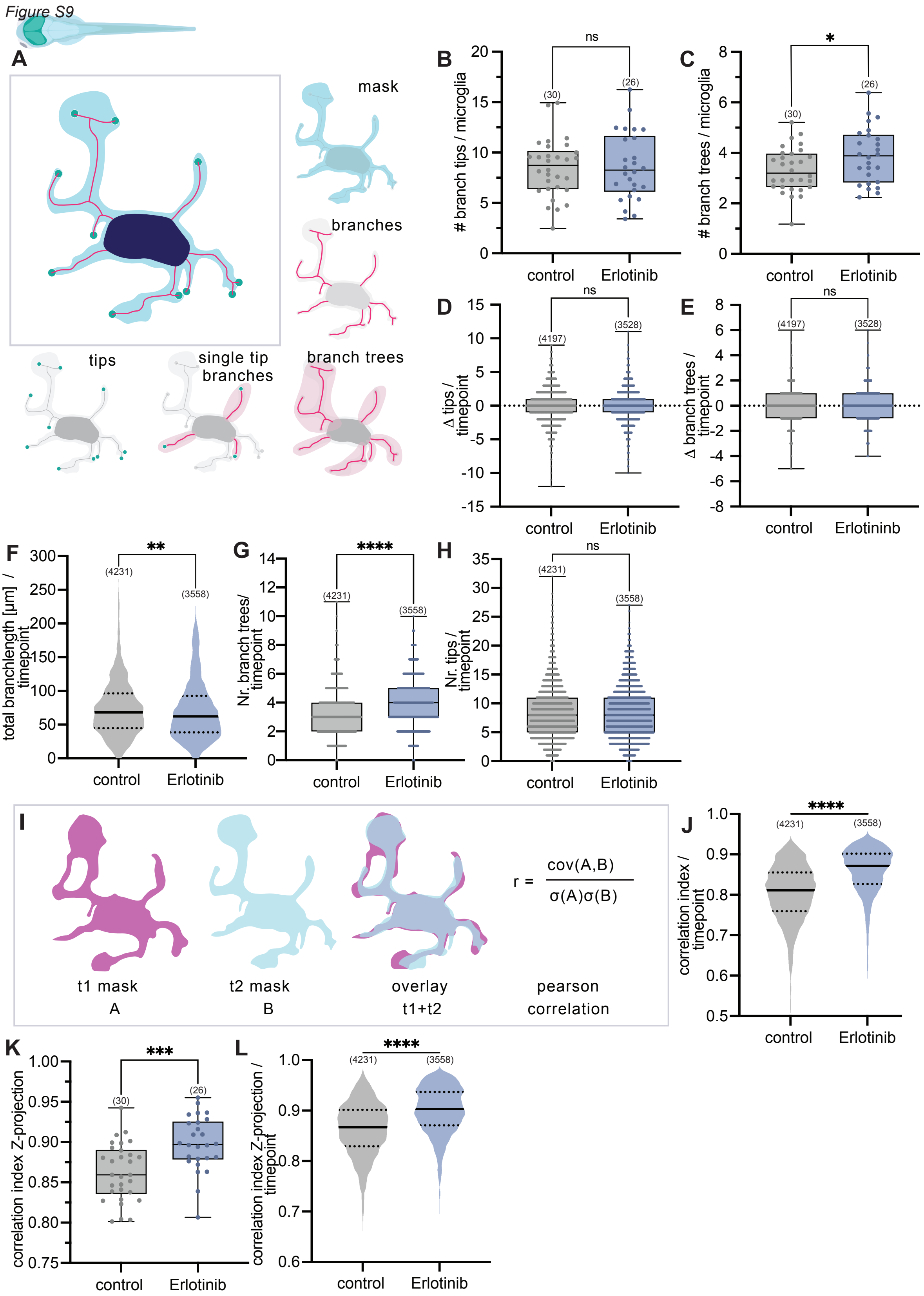

146

147 **Figure S9. Additional analysis of microglial mobility using per-time-point statistics.**

**(A)** Schematic overview of extracted microglial morphology features. Masks and branches were generated as illustrated in **Figure S10** and **Video S3**. Branch trees were defined as a connected skeleton with a single root (soma-connected component) and one or more branch tips defined as outer end points of the skeleton. Single-tip branches are defined as branches which contain only one tip. More information on the definition and creation of the structures can be found in Code Documentation **(Data S4)**. **(B)** Total number of branch tips per microglia over time, **(C)** total number of branch trees per microglia over time, **(D)** changes in the number of branch tips per time point, **(E)** changes in the number of branch trees per time point, **(F)** branch length per time point, **(G)** number of branch trees per time point, and **(H)** number of branch tips per time point analyses in control and Erlotinib-treated zebrafish. **(I)** Schematic illustration and formula used for the calculation of the Pearson correlation coefficient between two consecutive time points. ”A” corresponds to the mask at time point 1, and “B” to time point 2. Cov(A,B) represents the covariance and $\sigma$ the standard deviation between the two masks**.** A higher Pearson correlation coefficient indicates stronger mask overlap (i.e., lower mobility). **(J-L)** Additional calculations of the Pearson correlation coefficient. **(J)** **I**ndividual correlation coefficients between consecutive time points calculated from 3D binary masks, **(K)** mean correlation coefficients per microglial cell calculated from maximum intensity-projections of the masks over 60 min of recording, and **(L)** individual correlation coefficients between consecutive time points calculated from maximum intensity-projected masks. Numbers in brackets in **(B,C,K)** refer to the number of cells analyzed, in **(D,E)** to the number of consecutive timepoints and in **(F-H,J,L)** to the total number of timepoints analyzed. Boxplots show the median and interquartile range, whiskers indicate the minimum and maximum values. Violin plots with smoothing show the median (black line) and quartiles (dotted lines). Statistical tests used: Unpaired, two-tailed Welch’s tests. Statistical significance is indicated as ns ($p \geq 0.05$), * ($p < 0.05$), ** ($p < 0.01$), *** ($p < 0.001$), **** ($p < 0.0001$).

*Figure S10*

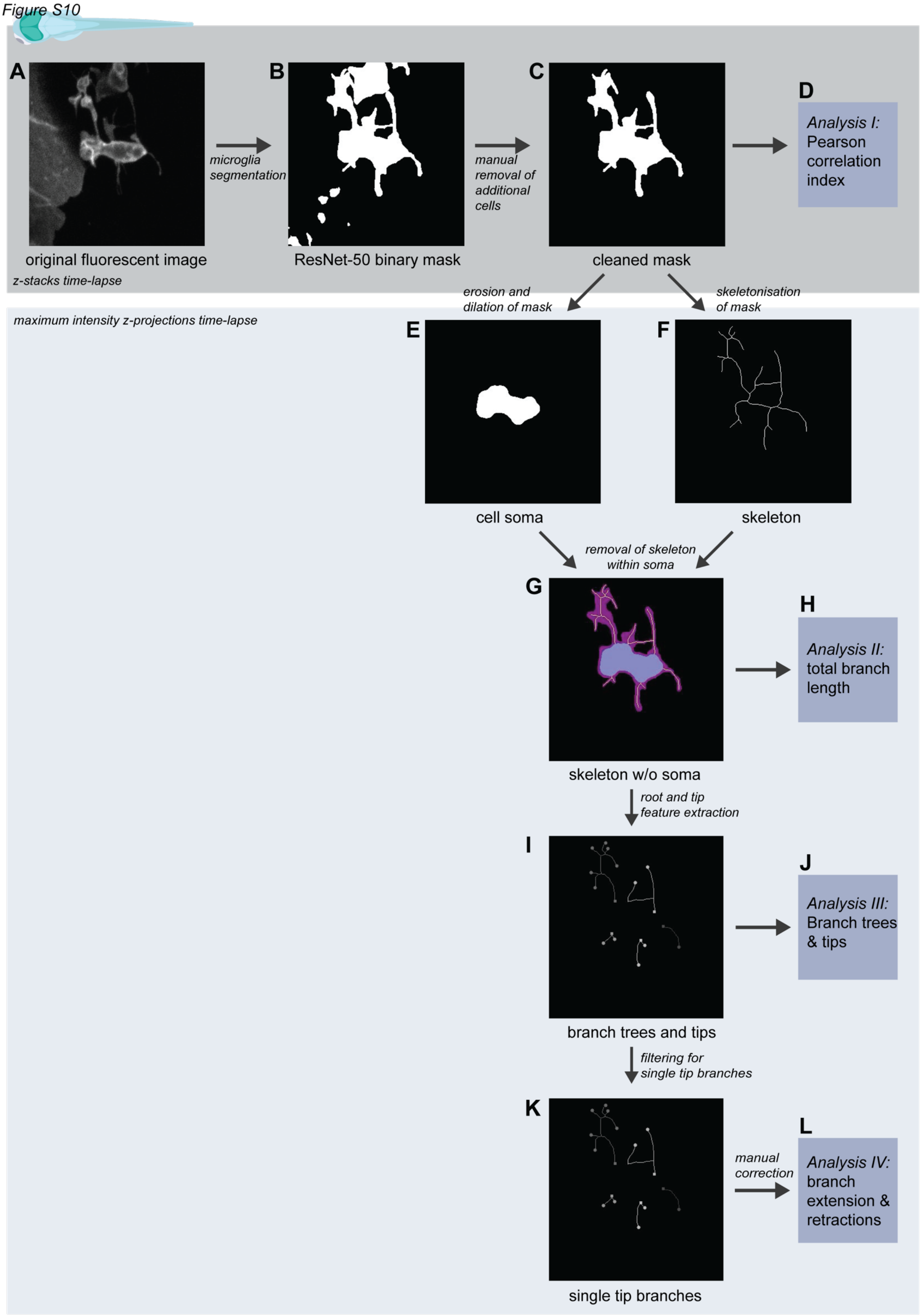
A
B
C
D
microglia segmentation
manual removal of additional cells
Analysis I: Pearson correlation index
original fluorescent image
ResNet-50 binary mask
cleaned mask
z-stacks time-lapse
erosion and dilation of mask
skeletonisation of mask
maximum intensity z-projections time-lapse
E
F
cell soma
skeleton
removal of skeleton within soma
G
H
Analysis II: total branch length
skeleton w/o soma
root and tip feature extraction
I
J
Analysis III: Branch trees & tips
branch trees and tips
filtering for single tip branches
K
L
manual correction
Analysis IV: branch extension & retractions
single tip branches

**Figure S10. Workflow of microglial segmentation and mobility analysis.**

See **Data S4** (Code Documentation) for the scripts used at each step. Scripts are named according to the panels **A-J** shown in this figure. **Video S3** provides an animated version of this figure. A maximum-intensity projection of the same control microglia is shown at each step. Each image stack was cropped to contain a single microglial cell before analysis. Microglial cells were imaged for at least 1 h at 30 s intervals. (A) Maximum-intensity projection of the original fluorescent image, **(B)** raw binary mask created by semantic segmentation, **(C)** cleaned mask (Masks were manually cleaned to remove masks of secondary microglia or skin cells. No masks were manually added.), **(D)** analysis of microglial motility by calculating the Pearson correlation coefficients **(Figure 6G and Figure S9I-L)**, **(E)** generation of the soma mask by removing branches (Temporal overlap, area stability, and centroid displacement constraints were applied to prevent transient phagosomes from being misclassified as soma.), **(F)** skeletonized mask representing the microglial structure, **(G)** skeleton after subtraction of the soma mask representing the branch structure, **(H)** analysis of total branch length **(Figure 6F and Figure S9F)**, **(I)** detection of branch trees and tips. A branch tree was defined as a connected skeleton component with a root connected to the soma mask. Each branch tree contains a single root and one or more branch tips. Branch tips were defined as endpoints of the skeleton excluding endpoints of short internal gaps that were merged to a single branch tree., **(J)** analysis of branch trees and tip numbers over time **(Figure S9B-E,G,H)**, **(K)** labeling of single tip branches. Single tip branches were defined as branches containing one root and one tip across multiple time points. The root was used for tracking the branches. Single-tip branches were tracked for at least two consecutive time points. Tracks were manually verified, and incorrect tracks were manually removed. No additional tracks were manually added, and **(L)** analysis of branch extension and retraction speed **(Figure 6H)**. Length changes between consecutive frames were classified as extension (increase in branch length) or retraction (decrease in branch length).

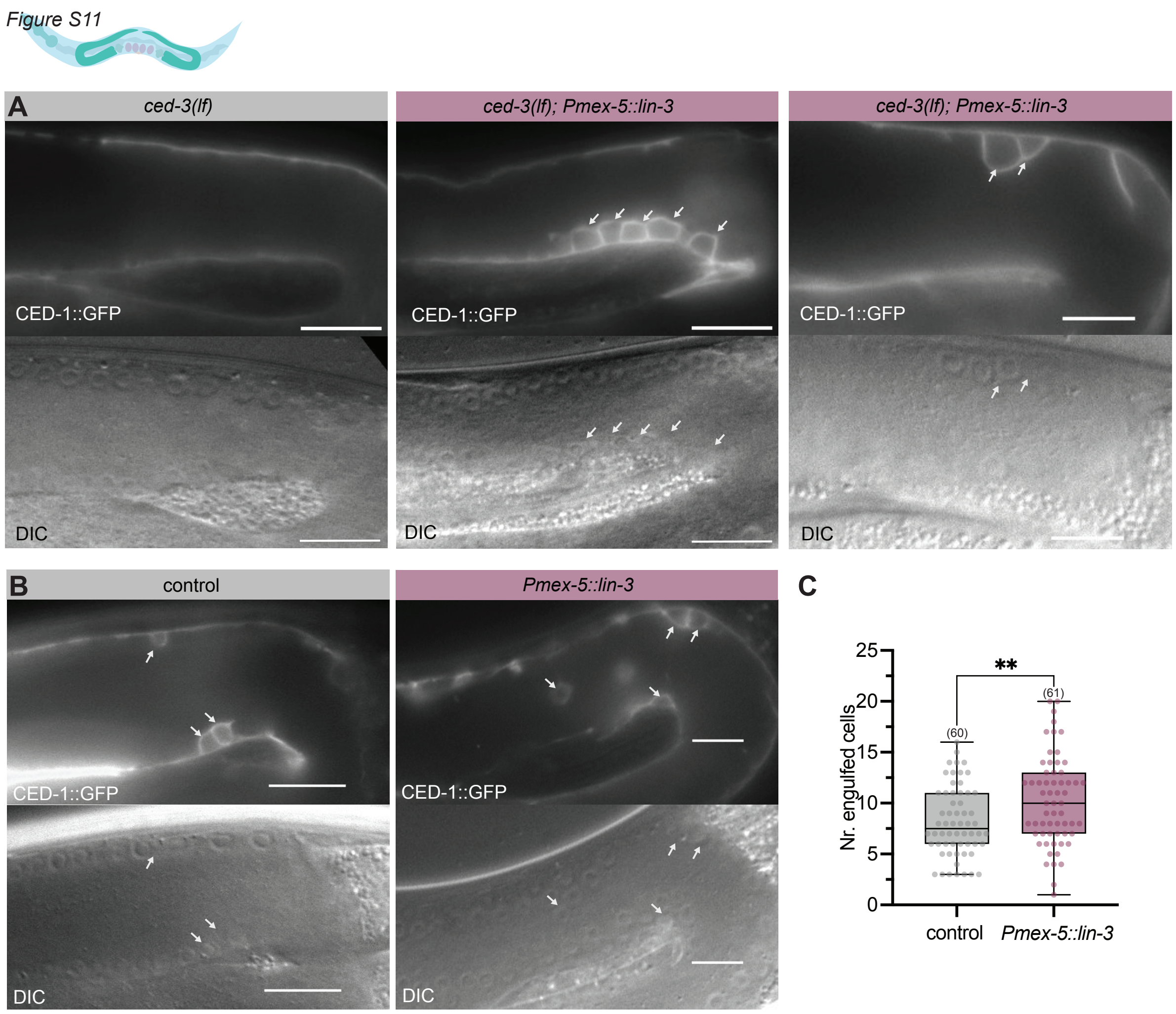

**Figure S11. CED-1::GFP-positive germ cells in *Pmex-5::lin-3* animals and quantification of** **engulfment in a wild-type background.**

**(A)** CED-1::GFP expression and corresponding DIC images of *Pmex-5::lin-3* animals in the apoptosis-deficient *ced-3(lf)* background. **(B)** CED-1::GFP expression and corresponding DIC images of *Pmex-* *5::lin-3* animals in a wild-type background. Single z-slices are shown, and arrows indicate engulfed cells. **(C)** Quantification of number of engulfed cells in *Pmex-5::lin-3* animals in a wild-type background. Boxplots show the median and interquartile range, whiskers indicate the minimum and maximum values. Statistical test used in **(C)**: Unpaired, two-tailed Welch's test. Statistical significance is indicated as ns ($p \geq 0.05$), * ($p < 0.05$), ** ($p < 0.01$), *** ($p < 0.001$), **** ($p < 0.0001$). Scale bars: 20 μm.
