## Supplementary material for "Conserved role of EGFR signaling in apoptotic cell recognition and processing across different phagocytes": Data S4

### Data S4: Code Documentation

#### General notes

*For the scripts made for this study, we in part used ChatGPT (OpenAI, GPT-5.2) during code development. Here, we list the scripts we created for the workflows, organised by the figure in which the resulting data was presented. During the code development, we additionally created intermediary scripts for parameter selection using visual outputs which are not listed here.*

*This document serves as a guide through the data analysis and references the scripts and where they were used.*

*Information on the code input format, dependencies, the goal of the script and the output created by the script is written in the code file header.*

#### Software versions

- Matlab R2023b (The MathWorks, Inc., Natick, MA, USA)
- Python 3.10.19

*All packages incl. the used versions installed in python environment are listed at the end of the document.*

---

#### Figure 2H,I, Supplementary Figure 2B-D, S3B,C

##### Flatfield correction for modERK-KTR intensity measurements

Prior to intensity measurements of the modERK-KTR sensor, a flat-field correction was performed using a custom MATLAB script. The correction normalizes illumination inhomogeneity using images of the illumination distribution for each channel and camera noise. The script generates flat-field corrected hyperstacks for downstream analysis. The flat-field correction code is not included in the code list.

$$Corrected_{Image} = m \cdot \frac{Raw_{Image} - Illumination_{Image}}{Illumination_{Image} - Noise_{Image}}$$

$$m = average(Illumination_{Image} - Noise_{Image})$$

---

#### *C. elegans* representative images across figures

##### Deconvolution

Across the figures containing *C. elegans* representative images (excluding the modErk-KTR images), we performed image deconvolution using a custom MATLAB script implementing Richardson-Lucy deconvolution[1] .Synthetic PSFs were generated using a Gibson-Lanni optical model[2]. The deconvolution script code is not included in the code list.

---

#### Figure 6C

##### Microglial migration upon EGF injection

To create the vector plot in Figure 6D, we used the following script:

6C_EGF_injection_vectorplot.py

The input data must be an excel file in this format:

| Track | X_start | Y_start | X_end | Y_end |
|---|---|---|---|---|
| 1 | 115.7544 | -82.0075 | 121.279 | -88.454 |
| 2 | 86.9104 | -24.4716 | 49.88809 | -5.57121 |

…

#### Figure 6E-I, Supplementary Figures S8-S10

#### Microglia mobility analysis

For the segmentation of microglia to the branch dynamics, we used a workflow with several different scripts. Please reference the **Supplementary Figure 10** for the workflow here described. The code files are named after the figure panels corresponding to the step in which the script was used.

To quantify microglia mobility, we started from time-lapse z-stacks of single microglia which were manually cropped from movies of living zebrafish larvae at 3 days post fertilization (3dpf). This approach facilitates the feature extraction per microglia and resulted in small 4D stacks (average of 1GB per microglia time track). The cropped single microglia vary in XY dimensions but have the same Z range. As a result, this pipeline allows for different XY dimensions. To use this pipeline with variable Z ranges, the according parts of the scripts would have to be adjusted.

To ensure labelling of the branches including fine branch structures, we used the fms:lyn-miRFP670 reporter line, which has suitable membrane labelling.

##### **S10B:** creating 4D microglia binary masks using a neural network

For the initial segmentation of the microglia to binary masks, we used a neural network approach. All subsequent steps after the initial mask creation are done without neural networks.

To generate the training data set, we made a randomization script, which selected random single Z slices across both the control and Erlotinib condition and across time. After creating the training dataset, we cropped the annotated slices to 256x256 pixels for training. As most stacks are significantly larger than 256x256, we cropped the annotated image and mask pair and generated five matched 256×256 training tiles per slice by cropping the pair at the four corners (top-left, top-right, bottom-left, bottom-right) and the centre. Additionally, we removed empty slices not containing a mask, as we have found that empty slices make to model worse and not better in our hands. In the end, with a training set of 2111 mask-image-pairs originating from 417 individual slices.

Using the annotated slices, we trained a model to automatically segment the microglia stacks. The segmentation model was based on a ResNet architecture[3] implemented in MATLAB. The trained model:

S10B_resnet50_1000.mat

For the first step of creating the binary masks, we used the MATLAB software.

The input consisted of the single microglia cropped 4D stacks (x,y,z,t). containing only the lyn-miRFP670 channel.

---

S10B_Segment_Microglia_4D.m

---

The output contained the segmented binary masks of the target microglia in the cropped region but occasionally it also contained parts of secondary microglia and of skin cells (which are also marked in fms:lyn-miRFP670).

**S10C**: Manual cleanup of 4D microglia masks

From the segmentation, there were often additional cells in the cropped stack of the microglia. To ensure we only have the mask of a single microglia per cropped stack, we manually cleaned the masks in Napari[4] . Undesired regions were erased by painting label 0 (background); no masks were manually added.

---

S10C_manual_microglial_mask_correction.py

---

After manual correction, a second script was used to remove small residual mask fragments:

---

S10C_2_remove_small_components.py

---

**S10D**: Pearson correlation of masks over time

Using the corrected masks, we then ran a script calculating the Pearson correlation coefficient of the masks between consecutive timepoints.

---

S10D_pearson_correlation_Zstack_and_Zproj_masks.py

---

As an input, we script accepts both the masks of a Z-stack over time or max Z-projection (produced in S6E_zproj_and_skeleton.py) over time. This script gives 4 different outputs organised in 2 separate excel files, “pearson_results_4D.xlsx” (Pearson correlation across 3D voxels over time) & “pearson_results_Z.xlsx” (Pearson correlation across 2D pixels over time). Two sheets are created per excel file, where “timepoint_metrics” gives the result of each 2 consecutive timepoints. The “summary_per_movie” sheet gives the average Pearson correlation index per movie. The mask size is also included as a control.

**S10F**: Z-projection and skeletonization

From the cleaned up binary masks, we created max Z-projections of the microglia and performed subsequent branch analysis on the 2D projections. On the 2D masks, we performed skeletonization[5] . In our data, 3D skeletonization produced duplicated representations of the same branch across Z planes, thus we did only proceed with 2D masks.

---

S10F_zproj_and_skeleton.py

---

#### **S10E**: Cell soma mask creation

Skeletonization produces a single-pixel-wide structure that also includes signal within the soma. To define branches, we generated a soma mask to exclude the skeleton within the cell soma. To create the soma mask, we eroded the binary mask to remove branches, followed by smoothing and dilation to reconstruct the central cell body. Temporal overlap, area stability, and centroid displacement constraints were applied to prevent transient phagosomes from being misclassified as soma.

---

S10E_cell_soma_mask_creation.py

---

#### **S10G**: Removal of skeleton within the soma

Using the cell soma mask, we removed all skeleton pixels within the cell soma which results in a skeleton representing only the microglia branches.

---

S10G_skeleton_from_cell_soma_removal.py

---

#### **S10I**: Generation of branch tree identities and tracking

For downstream analyses (tip number, branch tree numbers, branch extension and retractions) we assigned branch trees identities. Distance transforms and spatial computations were implemented using SciPy[6]. A branch tree was defined as a connected skeleton component with a root defined near or at the soma (soma-connected component), plus orphan components that were merged according to gap-distance and orientation criteria (see script header for parameters). A branch tree contains a single root and can contain one or more branch tips. Branch tips were defined as endpoints of the skeleton excluding endpoints of internal short gaps which were merged to a single branch tree.

---

S10I_1_branch_tree_identities.py

---

**S10H.J**: Branch length, number of tips and number of branch trees feature extraction

The branch length was defined as the total number of pixels of the branch skeleton in pixels and um. For the number of tips and branch trees, we used the arguments as described in S10I. Additionally, a visual output is created as a “RGB validation TIFF”, which shows the skeleton, the valid branch IDs, branch tips (circles) and roots (squares). Valid branches are shown in color, and have a branch ID number next to them. Excluded branches (one or more arguments as described in the code header are not met) are shown in grey.

---

S10H_J_branch_feature_extraction_visual_output.py

---

In a second script, we summarized the features per movie:

---

S10H_J_2_feature_summary_per_movie.py

---

**S10K**: Tracking of single tip branch trees for tip extension and retraction

To measure tip extension and retraction, we selected single tip branches (no ramification) and measured changes in branch length between consecutive timepoints. To ensure correct matching of branch identity over time, root pixels were used for tracking. This approach was chosen because roots are more stable for tracking than tips, and we could not identify a reliable tip-matching approach without neural networks. Extension was defined as a positive length change between consecutive timepoints; retraction was defined as a negative change.

First, a file was created containing all the single tip branches, including their branch ID, branch length and delta branch length in pixels:

---

S10K_1_extract_single_tip_branches.py

---

As not all branch roots were consistently tracked across timepoints, we implemented a manual validation step cross-referencing the RGB validation output (from S6H_J_branch_feature_extraction_visual_output.py displaying branch IDs) to remove incorrectly tracked branches in the excel sheet created (from S6K_1_extract_single_tip_branches.py).

Lastly, we then used a script to recompute single tip branch extension and retraction from the corrected list and convert the skeleton pixel to um.

---

S10K_2_correct_single_tip_branches.py

---

### **Python environment:** all installed packages and versions

```yaml
channels:
  - conda-forge
  - defaults
dependencies:
  - _openmp_mutex=4.5=2_gnu
  - aiohappyeyeballs=2.6.1=pyhd8ed1ab_0
  - aiohttp=3.13.2=py310h535af70_0
  - aiosignal=1.4.0=pyhd8ed1ab_0
  - alabaster=1.0.0=pyhd8ed1ab_1
  - annotated-types=0.7.0=pyhd8ed1ab_1
  - app-model=0.4.0=pyhd8ed1ab_0
  - appdirs=1.4.4=pyhd8ed1ab_1
  - arrow=1.4.0=pyhcf101f3_0
  - arrow-cpp=21.0.0=h6358905_1
  - asciitree=0.3.3=py_2
  - astroid=3.3.11=py310h5588dad_1
  - asttokens=3.0.1=pyhd8ed1ab_0
  - async-timeout=5.0.1=pyhd8ed1ab_1
  - asyncssh=2.21.1=pyhd8ed1ab_0
  - atomicwrites=1.4.1=pyhd8ed1ab_1
  - attrs=25.4.0=pyh71513ae_0
  - autopep8=2.0.4=pyhd8ed1ab_0
  - aws-c-auth=0.9.0=h02ab6af_2
  - aws-c-cal=0.9.2=h02ab6af_1
  - aws-c-common=0.12.4=h02ab6af_0
  - aws-c-compression=0.3.1=h02ab6af_2
  - aws-c-event-stream=0.5.6=h02ab6af_0
  - aws-c-http=0.10.4=h02ab6af_0
  - aws-c-io=0.21.4=h02ab6af_0
  - aws-c-mqtt=0.13.3=h02ab6af_0
  - aws-c-s3=0.8.7=h02ab6af_0
  - aws-c-sdkutils=0.2.4=h02ab6af_1
  - aws-checksums=0.2.7=h02ab6af_1
  - aws-crt-cpp=0.34.0=h885b0b7_0
  - aws-sdk-cpp=1.11.638=hf0af688_0
  - babel=2.17.0=pyhd8ed1ab_0
  - backports=1.0=pyhd8ed1ab_5
  - backports.tarfile=1.2.0=pyhd8ed1ab_1
  - bcrypt=5.0.0=py310h034784e_1
  - beautifulsoup4=4.14.2=pyha770c72_0
  - bermuda=0.1.6=py310hf13f778_1
  - binaryornot=0.4.4=pyhd8ed1ab_2
  - black=25.1.0=pyha5154f8_0
  - bleach=6.2.0=pyh29332c3_4
  - bleach-with-css=6.2.0=h82add2a_4
  - bokeh=3.8.0=py310haa95532_0
  - brotli-python=1.2.0=py310hd3493e3_0
  - bzip2=1.0.8=h2bbff1b_6
  - c-ares=1.34.5=h731ff69_0
  - ca-certificates=2025.12.2=haa95532_0
  - cachey=0.2.1=pyh9f0ad1d_0
  - cattrs=25.3.0=pyhd8ed1ab_0
  - certifi=2025.11.12=pyhd8ed1ab_0
  - cffi=2.0.0=py310h29418f3_1
  - chardet=5.2.0=pyhd8ed1ab_3
  - charset-normalizer=3.4.4=pyhd8ed1ab_0
  - click=8.3.1=pyh7428d3b_0
  - cloudpickle=3.1.2=pyhd8ed1ab_0
  - colorama=0.4.6=pyhd8ed1ab_1
  - comm=0.2.3=pyhe01879c_0
  - contourpy=1.3.1=py310h214f63a_0
  - cookiecutter=2.6.0=pyhd8ed1ab_1
  - cryptography=46.0.3=py310he482ccc_0
  - cycler=0.11.0=pyhd3eb1b0_0
  - cytoolz=1.1.0=py310h29418f3_1
  - dask=2025.11.0=py310haa95532_0
  - dask-core=2025.11.0=pyhcf101f3_0
  - debugpy=1.8.17=py310h699e580_0
  - decorator=5.2.1=pyhd8ed1ab_0
  - defusedxml=0.7.1=pyhd8ed1ab_0
  - diff-match-patch=20241021=pyhd8ed1ab_1
  - dill=0.4.0=pyhd8ed1ab_0
  - distributed=2025.11.0=py310haa95532_0
  - docstring-to-markdown=0.17=pyhe01879c_0
  - docstring_parser=0.17.0=pyhd8ed1ab_0
  - docutils=0.21.2=pyhd8ed1ab_1
  - et_xmlfile=2.0.0=py310haa95532_0
  - exceptiongroup=1.3.0=pyhd8ed1ab_0
  - executing=2.2.1=pyhd8ed1ab_0
  - expat=2.7.3=h9214b88_0
  - fasteners=0.19=pyhd8ed1ab_1
  - flake8=7.1.2=pyhd8ed1ab_0
  - flexcache=0.3=pyhd8ed1ab_1
  - flexparser=0.4=pyhd8ed1ab_1
  - fonttools=4.61.0=py310h02ab6af_0
  - freetype=2.14.1=h57928b3_0
  - freetype-py=2.5.1=pyhd8ed1ab_1
  - frozenlist=1.7.0=py310had1666a_0
  - fsspec=2025.10.0=pyhd8ed1ab_0
  - fzf=0.67.0=hb13f615_0
  - gflags=2.2.2=hd77b12b_1
  - glib=2.84.4=h60240c0_0
  - glib-tools=2.84.4=h3823947_0
  - glog=0.5.0=hd77b12b_1
  - gst-plugins-base=1.24.11=h3fe0a9e_0
  - gstreamer=1.24.11=h233a61a_0
  - h2=4.3.0=pyhcf101f3_0
  - heapdict=1.0.1=pyhd8ed1ab_2
  - hpack=4.1.0=pyhd8ed1ab_0
  - hsluv=5.0.4=pyhd8ed1ab_2
  - hyperframe=6.1.0=pyhd8ed1ab_0
  - icu=75.1=he0c23c2_0
  - idna=3.11=pyhd8ed1ab_0
  - imageio=2.37.0=pyhfb79c49_0
  - imagesize=1.4.1=pyhd8ed1ab_0
  - importlib-metadata=8.7.0=pyhe01879c_1
  - importlib_resources=6.5.2=pyhd8ed1ab_0
  - in-n-out=0.2.1=pyhd8ed1ab_1
  - inflection=0.5.1=pyhd8ed1ab_1
  - intervaltree=3.1.0=pyhd8ed1ab_1
  - ipykernel=6.31.0=pyh6dadd2b_0
  - ipython=8.37.0=pyha7b4d00_0
  - ipython_pygments_lexers=1.1.1=pyhd8ed1ab_0
  - isort=6.1.0=pyhd8ed1ab_0
  - jaraco.classes=3.4.0=pyhd8ed1ab_2
  - jaraco.context=6.0.1=pyhd8ed1ab_0
  - jaraco.functools=4.3.0=pyhd8ed1ab_0
  - jedi=0.19.2=pyhd8ed1ab_1
  - jellyfish=1.1.3=py310hc226416_0
  - jinja2=3.1.6=pyhd8ed1ab_0
  - joblib=1.5.2=pyhd8ed1ab_0
  - jsonschema=4.25.1=pyhe01879c_0
  - jsonschema-specifications=2025.9.1=pyhcf101f3_0
  - jupyter_client=8.6.3=pyhd8ed1ab_1
  - jupyter_core=5.9.1=pyh6dadd2b_0
  - jupyterlab_pygments=0.3.0=pyhd8ed1ab_2
  - keyring=25.7.0=pyh7428d3b_0
  - kiwisolver=1.4.9=py310h1e1005b_2
  - krb5=1.21.3=hdf4eb48_0
  - lazy-loader=0.4=pyhd8ed1ab_2
  - lazy_loader=0.4=pyhd8ed1ab_2
  - lcms2=2.17=hbcf6048_0
  - lerc=4.0.0=h6470a55_1
  - libabseil=20250127.0=cxx17_h52369b4_0
  - libblas=3.9.0=39_hf2e6a31_mkl
  - libbrotlicommon=1.0.9=h827c3e9_9
  - libbrotlidec=1.0.9=h827c3e9_9
  - libbrotlienc=1.0.9=h827c3e9_9
  - libcblas=3.9.0=39_h2a3cdd5_mkl
  - libclang13=21.1.5=default_ha2db4b5_0
  - libcurl=8.16.0=h97e0424_0
  - libdeflate=1.22=h2466b09_0
  - libffi=3.4.4=hd77b12b_1
  - libfreetype=2.14.1=h57928b3_0
  - libfreetype6=2.14.1=hdbac1cb_0
  - libgcc=15.2.0=h1383e82_7
  - libglib=2.84.4=hfaec014_0
  - libgomp=15.2.0=h1383e82_7
  - libgrpc=1.71.0=hf4237ab_0
  - libhwloc=2.12.1=default_h88281d1_1000
  - libiconv=1.18=hc1393d2_2
  - libintl=0.22.5=h5728263_3
  - libjpeg-turbo=3.1.2=hfd05255_0
  - liblapack=3.9.0=39_hf9ab0e9_mkl
  - libogg=1.3.5=h2466b09_1
  - libpng=1.6.50=h7351971_1
  - libprotobuf=5.29.3=h65a231f_1
  - libre2-11=2024.07.02=h5da7b33_0
  - libsodium=1.0.20=hc70643c_0
  - libspatialindex=2.1.0=h518811d_0
  - libsqlite=3.51.0=hf5d6505_0
  - libssh2=1.11.1=h2addb87_0
  - libthrift=0.22.0=ha2884a9_0
  - libtiff=4.7.0=hfc51747_1
  - libvorbis=1.3.7=h5112557_2
  - libwebp=1.6.0=h4d5522a_0
  - libwebp-base=1.6.0=h4d5522a_0
  - libwinpthread=12.0.0.r4.gg4f2fc60ca=h57928b3_10
  - libxcb=1.17.0=h0e4246c_0
  - libxml2=2.13.9=h741aa76_0
  - libzlib=1.3.1=h02ab6af_0
  - llvm-openmp=21.1.5=h4fa8253_2
  - llvmlite=0.45.1=py310hfe4b161_0
  - locket=1.0.0=pyhd8ed1ab_0
  - lsprotocol=2025.0.0=pyhe01879c_0
  - lz4=4.4.5=py310hb1efef3_0
  - lz4-c=1.9.4=hcfcfb64_0
  - magicgui=0.10.1=pyhd8ed1ab_0
  - markdown-it-py=4.0.0=pyhd8ed1ab_0
  - markupsafe=3.0.3=py310hdb0e946_0
  - matplotlib=3.10.7=py310haa95532_0
  - matplotlib-base=3.10.7=py310h26e45b9_0
  - matplotlib-inline=0.2.1=pyhd8ed1ab_0
  - mccabe=0.7.0=pyhd8ed1ab_1
  - mdurl=0.1.2=pyhd8ed1ab_1
  - mistune=3.1.4=pyhcf101f3_0
  - mkl=2025.3.0=hac47afa_454
  - more-itertools=10.8.0=pyhd8ed1ab_0
  - msgpack-python=1.1.2=py310he9f1925_1
  - multidict=6.6.3=py310hdb0e946_0
  - mypy_extensions=1.1.0=pyha770c72_0
  - napari=0.6.6=pyh4da3b27_0
  - napari-base=0.6.6=pyhcf101f3_0
  - napari-console=0.1.4=pyhc04f84c_0
  - napari-plugin-engine=0.2.0=pyha07c04f_3
  - napari-plugin-manager=0.1.8=pyha44d970_0
  - napari-svg=0.2.1=pyha07c04f_0
  - narwhals=2.7.0=py310haa95532_0
  - nbclient=0.10.2=pyhd8ed1ab_0
  - nbconvert=7.16.6=hc388f54_1
  - nbconvert-core=7.16.6=pyhcf101f3_1
  - nbconvert-pandoc=7.16.6=h7d6f222_1
  - nbformat=5.10.4=pyhd8ed1ab_1
  - nest-asyncio=1.6.0=pyhd8ed1ab_1
  - networkx=3.4.2=pyh267e887_2
  - npe2=0.7.9=pyhd8ed1ab_0
  - numba=0.62.1=py310h86ba7b5_0
  - numcodecs=0.13.1=py310hb4db72f_0
  - numpy=2.2.6=py310h4987827_0
  - numpydoc=1.9.0=pyhe01879c_1
  - openjpeg=2.5.3=h24db6dd_1
  - openpyxl=3.1.5=py310h827c3e9_1
  - openssl=3.6.0=h725018a_0
  - orc=2.2.0=hf43e1a1_0
  - packaging=25.0=pyh29332c3_1
  - pandas=2.3.3=py310hed136d8_1
  - pandoc=3.8.2.1=h57928b3_0
  - pandocfilters=1.5.0=pyhd8ed1ab_0
  - parso=0.8.5=pyhcf101f3_0
  - partd=1.4.2=pyhd8ed1ab_0
  - partsegcore-compiled-backend=0.15.14=py310hed136d8_1
  - pathspec=0.12.1=pyhd8ed1ab_1
  - pcre2=10.46=h3402e2f_0
  - pexpect=4.9.0=pyhd8ed1ab_1
  - pickleshare=0.7.5=pyhd8ed1ab_1004
  - pillow=11.3.0=py310hb3a2f59_3
  - pint=0.24.4=pyhe01879c_2
  - pip=25.3=pyhc872135_0
  - platformdirs=4.5.0=pyhcf101f3_0
  - pluggy=1.6.0=pyhd8ed1ab_0
  - ply=3.11=pyhd8ed1ab_3
  - pooch=1.8.2=pyhd8ed1ab_3
  - prompt-toolkit=3.0.52=pyha770c72_0
  - propcache=0.3.1=py310h38315fa_0
  - psutil=7.1.3=py310h1637853_0
  - psygnal=0.15.0=pyhd8ed1ab_0
  - pthread-stubs=0.4=h0e40799_1002
  - ptyprocess=0.7.0=pyhd8ed1ab_1
  - pure_eval=0.2.3=pyhd8ed1ab_1
  - pyarrow=21.0.0=py310ha5e6156_0
  - pycodestyle=2.12.1=pyhd8ed1ab_1
  - pyconify=0.2.1=pyhd8ed1ab_0
  - pycparser=2.22=pyh29332c3_1
  - pydantic=2.12.4=pyh3cfb1c2_0
  - pydantic-compat=0.1.2=pyhd8ed1ab_0
  - pydantic-core=2.41.5=py310h034784e_1
  - pydocstyle=6.3.0=pyhd8ed1ab_1
  - pyflakes=3.2.0=pyhd8ed1ab_1
  - pygithub=2.8.1=py310haa95532_0
  - pygments=2.19.2=pyhd8ed1ab_0
  - pyjwt=2.10.1=pyhd8ed1ab_0
  - pylint=3.3.9=pyhcf101f3_0
  - pylint-venv=3.0.4=pyhd8ed1ab_1
  - pyls-spyder=0.4.0=pyhd8ed1ab_1
  - pynacl=1.6.0=py310h29418f3_0
  - pyopengl=3.1.10=pyh7428d3b_2

  - pyopenssl=25.3.0=pyhd8ed1ab_0
  - pyparsing=3.2.5=py310haa95532_0
  - pyproject_hooks=1.2.0=pyhd8ed1ab_1
  - pyqt=5.15.11=py310hdf200a9_2
  - pyqt5-sip=12.17.0=py310h73ae2b4_2
  - pyqtwebengine=5.15.11=py310h4f81690_2
  - pysocks=1.7.1=pyh09c184e_7
  - python=3.10.19=h981015d_0
  - python-build=1.3.0=pyhff2d567_0
  - python-dateutil=2.9.0.post0=pyhe01879c_2
  - python-fastjsonschema=2.21.2=pyhe01879c_0
  - python-gssapi=1.10.1=py310hc1833cf_1
  - python-lmdb=1.6.2=py310h5da7b33_0
  - python-lsp-black=2.0.0=pyhff2d567_1
  - python-lsp-jsonrpc=1.1.2=pyhff2d567_1
  - python-lsp-ruff=2.3.0=pyhcf101f3_0
  - python-lsp-server=1.13.1=pyh332efcf_0
  - python-lsp-server-base=1.13.1=pyhd8ed1ab_0
  - python-slugify=8.0.4=pyhd8ed1ab_1
  - python-tzdata=2025.2=pyhd8ed1ab_0
  - python_abi=3.10=2_cp310
  - pytoolconfig=1.2.5=pyhd8ed1ab_1
  - pytz=2025.2=pyhd8ed1ab_0
  - pyuca=1.2=pyhd8ed1ab_2
  - pywavelets=1.8.0=py310hb0944cc_0
  - pywin32=311=py310h282bd7d_1
  - pywin32-ctypes=0.2.3=py310h5588dad_3
  - pyyaml=6.0.3=py310hdb0e946_0
  - pyzmq=27.1.0=py310h535538e_0
  - qdarkstyle=3.2.3=pyhd8ed1ab_1
  - qstylizer=0.2.4=pyhff2d567_0
  - qt-main=5.15.15=hb098fff_5
  - qt-webengine=5.15.15=h087ee03_1
  - qtawesome=1.4.0=pyh9208f05_1
  - qtconsole=5.7.0=pyhd8ed1ab_0
  - qtconsole-base=5.7.0=pyha770c72_0
  - qtpy=2.4.3=pyhd8ed1ab_1
  - re2=2024.07.02=h214f63a_0
  - referencing=0.37.0=pyhcf101f3_0
  - requests=2.32.5=pyhd8ed1ab_0
  - rich=14.2.0=pyhcf101f3_0
  - rope=1.14.0=pyhd8ed1ab_0
  - rpds-py=0.29.0=py310h034784e_0
  - rtree=1.4.1=pyh11ca60a_0
  - ruff=0.14.5=h15e3a1f_0
  - scikit-image=0.25.2=py310hed136d8_2
  - scikit-learn=1.7.2=py310h21054b0_0
  - scipy=1.15.2=py310h15c175c_0
  - setuptools=80.9.0=py310haa95532_0
  - shellingham=1.5.4=pyhd8ed1ab_2
  - sip=6.10.0=py310h73ae2b4_1
  - six=1.17.0=pyhe01879c_1
  - snappy=1.2.2=h7fa0ca8_1
  - snowballstemmer=3.0.1=pyhd8ed1ab_0
  - sortedcontainers=2.4.0=pyhd8ed1ab_1
  - soupsieve=2.8=pyhd8ed1ab_0
  - sphinx=8.1.3=pyhd8ed1ab_1
  - sphinxcontrib-applehelp=2.0.0=pyhd8ed1ab_1
  - sphinxcontrib-devhelp=2.0.0=pyhd8ed1ab_1
  - sphinxcontrib-htmlhelp=2.1.0=pyhd8ed1ab_1
  - sphinxcontrib-jsmath=1.0.1=pyhd8ed1ab_1
  - sphinxcontrib-qthelp=2.0.0=pyhd8ed1ab_1
  - sphinxcontrib-serializinghtml=1.1.10=pyhd8ed1ab_1
  - spyder=6.1.0=hd8ed1ab_0
  - spyder-base=6.1.0=py310h5588dad_0
  - spyder-kernels=3.1.1=win_pyh7428d3b_0
  - sqlite=3.51.0=hda9a48d_0
  - stack_data=0.6.3=pyhd8ed1ab_1
  - superqt=0.7.6=pyhb6d5dde_0
  - tbb=2022.3.0=hd094cb3_1
  - tblib=3.1.0=py310haa95532_0
  - text-unidecode=1.3=pyhd8ed1ab_2
  - textdistance=4.6.3=pyhd8ed1ab_1
  - threadpoolctl=3.6.0=pyhecae5ae_0
  - three-merge=0.1.1=pyhd8ed1ab_1
  - tifffile=2025.2.18=py310haa95532_0
  - tinycss2=1.4.0=pyhd8ed1ab_0
  - tk=8.6.15=hf199647_0
  - toml=0.10.2=pyhd8ed1ab_2
  - tomli=2.3.0=pyhcf101f3_0
  - tomli-w=1.2.0=pyhd8ed1ab_0
  - tomlkit=0.13.3=pyha770c72_0
  - toolz=1.1.0=pyhd8ed1ab_1
  - tornado=6.5.2=py310h29418f3_2
  - tqdm=4.67.1=pyhd8ed1ab_1
  - traitlets=5.14.3=pyhd8ed1ab_1
  - typer=0.20.0=pyhefaf540_1
  - typer-slim=0.20.0=pyhcf101f3_1
  - typer-slim-standard=0.20.0=h4daf872_1
  - typing-extensions=4.15.0=h396c80c_0
  - typing-inspection=0.4.2=pyhd8ed1ab_0
  - typing_extensions=4.15.0=pyhcf101f3_0
  - tzdata=2025b=h04d1e81_0
  - ucrt=10.0.22621.0=haa95532_0
  - ujson=5.11.0=py310h699e580_1
  - urllib3=2.5.0=pyhd8ed1ab_0
  - utf8proc=2.6.1=h2bbff1b_1
  - vc=14.3=h2df5915_10
  - vc14_runtime=14.44.35208=h4927774_10
  - vispy=0.15.2=py310h8f3aa81_1
  - vs2015_runtime=14.44.35208=ha6b5a95_10
  - watchdog=6.0.0=py310h5588dad_2
  - wcwidth=0.2.14=pyhd8ed1ab_0
  - webencodings=0.5.1=pyhd8ed1ab_3
  - whatthepatch=1.0.7=pyhd8ed1ab_1
  - wheel=0.45.1=py310haa95532_0
  - win_inet_pton=1.1.0=pyh7428d3b_8
  - wrapt=2.0.1=py310h29418f3_1
  - xorg-libxau=1.0.12=hba3369d_1
  - xorg-libxdmcp=1.1.5=hba3369d_1
  - xyzservices=2025.4.0=py310haa95532_0
  - xz=5.6.4=h4754444_1
  - yaml=0.2.5=h6a83c73_3
  - yapf=0.43.0=pyhd8ed1ab_1
  - yarl=1.22.0=py310hdb0e946_0
  - zarr=2.18.3=pyhd8ed1ab_1
  - zeromq=4.3.5=h5bddc39_9
  - zict=3.0.0=py310haa95532_0
  - zipp=3.23.0=pyhd8ed1ab_0
  - zlib=1.3.1=h02ab6af_0
  - zstandard=0.25.0=py310h1637853_1
  - zstd=1.5.7=hbeecb71_2
  - pip:
    - h5py==3.15.1
    - imagecodecs==2025.3.30
    - imaris-ims-file-reader==0.1.8
    - opencv-python==4.12.0.88
```
